# Molecular phylogeny of fucoxanthin-chlorophyll *a/c* proteins from *Chaetoceros gracilis* and Lhcq/Lhcf diversity

**DOI:** 10.1101/2021.09.06.459020

**Authors:** Minoru Kumazawa, Hiroyo Nishide, Ryo Nagao, Natsuko Inoue-Kashino, Jian-Ren Shen, Takeshi Nakano, Ikuo Uchiyama, Yasuhiro Kashino, Kentaro Ifuku

## Abstract

Diatoms adapt to various aquatic light environments and play major roles in the global carbon cycle using their unique light-harvesting system, i.e., fucoxanthin chlorophyll *a*/*c* binding proteins (FCPs). Structural analyses of photosystem II (PSII)-FCPII and photosystem I (PSI)-FCPI complexes from the diatom *Chaetoceros gracilis* have revealed the localization and interactions of many FCPs; however, the entire set of FCPs has not been characterized. Here, we identified 46 FCPs in the newly assembled genome and transcriptome of *C. gracilis*. Phylogenetic analyses suggested that these FCPs could be classified into five subfamilies: Lhcr, Lhcf, Lhcx, Lhcz, and novel Lhcq, in addition to a distinct type of Lhcr, CgLhcr9. The FCPs in Lhcr, including CgLhcr9 and some Lhcqs, had orthologous proteins in other diatoms, particularly those found in the PSI-FCPI structure. By contrast, the Lhcf subfamily, some of which were found in the PSII-FCPII complex, seemed to be diversified in each diatom species, and the number of Lhcqs differed among species, indicating that their diversification may contribute to species-specific adaptations to light. Further phylogenetic analyses of FCPs/light-harvesting complex (LHC) proteins using genome data and assembled transcriptomes of other diatoms and microalgae in public databases suggest that our proposed classification of FCPs was common among various red-lineage algae derived from secondary endosymbiosis of red algae, including Haptophyta. These results provided insights into the loss and gain of FCP/LHC subfamilies during the evolutionary history of the red algal lineage.

**One sentence summary:** Phylogenetic analysis of fucoxanthin-chlorophyll *a*/*c* proteins in *C. gracilis* revealed five major subfamilies and one minor subfamily, providing insights into the diversification of light-harvesting systems in red algae.

## Introduction

Diatoms are a group of photosynthetic Stramenopiles (or Heterokonts), which are red-lineage secondary symbiotic algae with plastids derived from Rhodophyta (red algae). Diatoms are major primary producers in modern oceans (José et al., 2019). Unlike the green lineage, diatoms have a brown color owing to the presence of photosynthetic pigments different from those of the green lineage (e.g., chlorophyll [Chl] *a*, Chl *c*, fucoxanthin, and diadinoxanthin) in the light-harvesting pigment protein complex (LHC) surrounding their photosystems. This LHC in diatoms is called fucoxanthin Chl *a*/*c* binding protein (FCP), which absorbs light with blue-green wavelengths and thus captures more light in aqueous environments. In addition, LHC/FCP can function in non-photochemical quenching (NPQ), which dissipates the excitation energy of excessively absorbed light as heat (Niyogi and Truong 2013; Ruban 2018; Goss and Lepetit 2015; Wobbe et al. 2016; Giovagnetti and Ruban 2018). The core subunits of photosynthetic protein complexes are highly conserved among oxygenic photosynthetic organisms; however, in most eukaryotic photosynthetic organisms, the LHC shows diversified sequences and pigment compositions to adapt to the living environment (Büchel, 2015; Büchel, 2020).

The first X-ray crystal structure of photosystem II of cyanobacteria was reported at the atomic level (Umena et al., 2011), and the structures of photosystems in green-lineage plants have been resolved by both X-ray crystallography and cryo-electron microscopy (EM) (Mazor et al., 2015; Qin et al., 2015; Wei et al., 2016; Mazor et al., 2017; Su et al., 2017; Shen et al., 2019). In addition, structures of photosystems in red-lineage plants have also been reported; photosystem II (PSII) of Rhodophyta *Cyanidium caldarium* was resolved by X-ray crystal structural analysis (Ago et al., 2016), whereas photosystem I of Rhodophyta *Cyanidioschyzon merolae* was resolved by cryo-EM (Pi et al., 2018). The structures of the PSII-FCPII supercomplex of the centric diatom *Chaetoceros gracilis* was reported as the first photosystem structure in secondary symbiotic algae (Nagao *et al*., 2019; Pi *et al*., 2019), followed by that of PSI-FCPI of *Chaetoceros gracilis* (Nagao et al., 2020). The structure of *Chaetoceros gracilis* PSI-FCPI, which has an increased number of FCPs, has also been reported (Xu et al., 2020). Accordingly, the molecular phylogeny of diatom FCPs can be interpreted on the basis of structural information and mass spectrometric identification of FCPs in the complexes separated by sucrose density gradient or native polyacrylamide gel electrophoresis (PAGE).

In our report on diatom PSI-FCPI (Nagao *et al*. 2020), we argued that FCPs in the outer edge of PSI-FCPI should belong to a novel group, Lhcq, a phylogenetic group different from that of Lhcr, which is commonly found in red-algal PSI (Nagao et al., 2020). Hoffman *et al*., (2011) reported the LHC/FCP phylogeny using expressed sequence tags of red-lineage species and the genome sequences of the red alga *Cyanidioschyzon merolae*, pennate diatom *Phaeodactylum tricornutum*, and centric diatom *Thalassiosira pseudonana*.

Here, we performed a more comprehensive analysis of the draft genome of *Chaetoceros gracilis*, its FCP sequences, and the LHC/FCP sequences obtained from the genomes of other diatoms and algae in the red lineage. Overall, our results suggest that the diversified subfamilies of LHC/FCP, particularly those of Lhcf and Lhcq, had occurred in the common ancestral origin of red lineage algae, contributing to their high adaptability and prosperity in the ocean.

## Results

### Assembly, gene prediction, and genome completeness

Next-generation sequencing (NGS) data suggested that the estimated size of the *Chaetoceros gracilis* genome was 35.4 Mbp. The assembled draft nuclear genome contained 791 scaffolds and 3,408 contigs with an N50 of 180 kbp and GC content of 37.3% (**Fig. 1A, B**). In total, 15,484 genes were predicted as nuclear-coded genes by BRAKER2 (Hoff et al., 2016). The assembly included chloroplast scaffolds with gene prediction performed using DOGMA. Benchmarking Universal Single-Copy Orthologs (BUSCO) suggested that 96% of conserved single-copy genes of *Stramenopiles_odb10* were included in the predicted genes in the *Chaetoceros gracilis* nuclear genome (**Fig. 1C**), similar to values for *Thalassiosira pseudonana* (97%) and *Phaeodactylum tricornutum* (97%; **Supplemental Table S1**), indicating that our *Chaetoceros gracilis* draft genome had adequate coverage of essential genes. Using OrthoFinder, the predicted nuclear-coded genes in the *Chaetoceros gracilis* genome were classified into 7,320 orthogroups, including 5,563 orthogroups (77%) common with *Thalassiosira pseudonana* and 5,451 orthogroups (74.5%) common with *Phaeodactylum tricornutum* (**Fig. 1B**).

**Figure 1.**
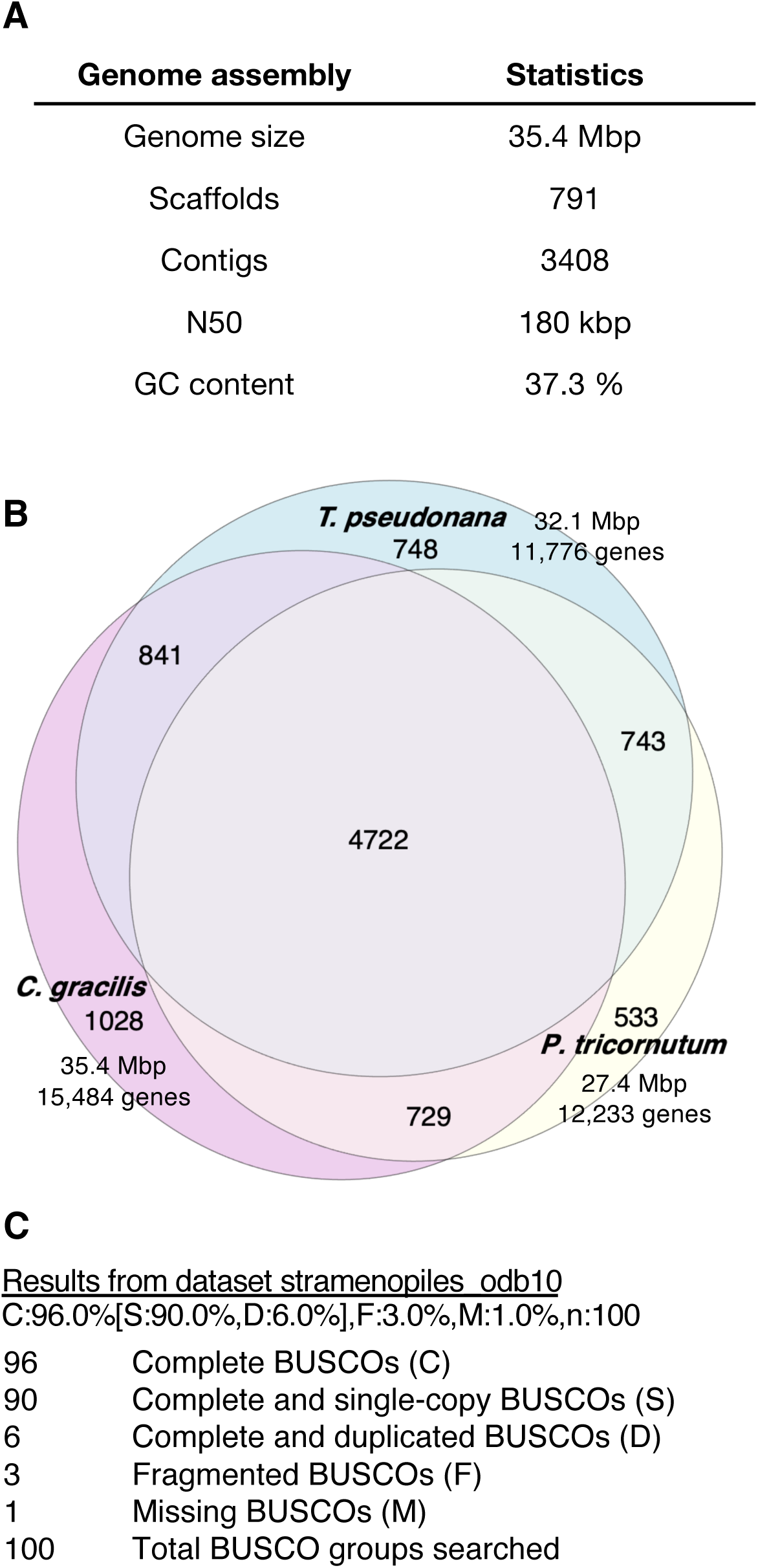
Assessments of the *Chaetoceros gracilis* draft genome assembly. **A**, General statistics of the *Chaetoceros gracilis* draft genome. **B**, Euler diagram of the orthogroups among *Chaetoceros gracilis* and two model diatom nuclear genomes, *Thalassiosira pseudonana* and *Phaeodactylum tricornutum*, with the draft genome size and the number of predicted genes. The diagram was generated using the Eulerr package (Wilkinson, 2012; Micallef and Rodgers, 2014) with R language. **C**, BUSCO scores for the predicted genes in the draft nuclear genome of *Chaetoceros gracilis* using the dataset Stramenopiles_odb10.

### Molecular phylogeny of FCPs obtained from genomes and transcriptomes

*Chaetoceros gracilis* FCPs (CgFCPs) were exhaustively searched using the draft genome and long-read transcriptome data. Forty-four CgFCPs were obtained from the *Chaetoceros gracilis* draft genome using the FCP genes of *Thalassiosira pseudonana* (TpFCPs, 30 genes) and *Phaeodactylum tricornutum* (PtFCPs, 39 genes) in the NCBI RefSeq database as queries. The CgFCPs were further complemented by a long-read transcriptome, IsoSeq, for *Chaetoceros gracilis* from two culture conditions. Transcriptomes were refined using IsoSeq3 and isONclust. Refined transcriptomes by IsoSeq3 had BUSCO scores of 73% and 78%, respectively, and those from isONclust had scores of 78% and 77%, respectively. Two CgFCPs, CgLhcf13 and CgLhcf14, which were not found in the draft genome, were detected by BLASTP search of transcriptomes using the same query set. Additionally, 44 TpFCPs and 42 PtFCPs were exhaustively extracted from RefSeq genomes (Armbrust et al., 2004; Bowler et al., 2008; Rastogi et al., 2018) using BLASTP similarity search with 30 TpFCPs, 39 PtFCPs, and 46 CgFCPs as a query set. This exhaustive FCP search revealed that *Chaetoceros gracilis, Thalassiosira pseudonana*, and *Phaeodactylum tricornutum* had 46, 44, and 42 FCPs, respectively (**Supplemental Tables S2–4**).

Phylogenetic analyses of CgFCPs with curated FCPs from *Thalassiosira pseudonana* (**Fig. 2A**) or *Phaeodactylum tricornutum* (**Fig. 2B**) suggested that CgFCPs could be systemically named using the four major types: Lhcr, Lhcf, and Lhcx annotated in previous studies (Koziol *et al*., 2007; Dittami *et al*., 2010; Hoffman *et al*., 2011) and the new subfamily named Lhcq. The Lhcr type included CgLhcr4 and CgLhcr9, as well as the Lhcz subfamily. Although CgLhcr4 and CgLhcr9 were not branched into the Lhcr clade in our phylogenetic analysis, they were included in “Lhcr” because of their locations in the PSI-FCPI complex (Nagao *et al*., 2020).

**Figure 2.**
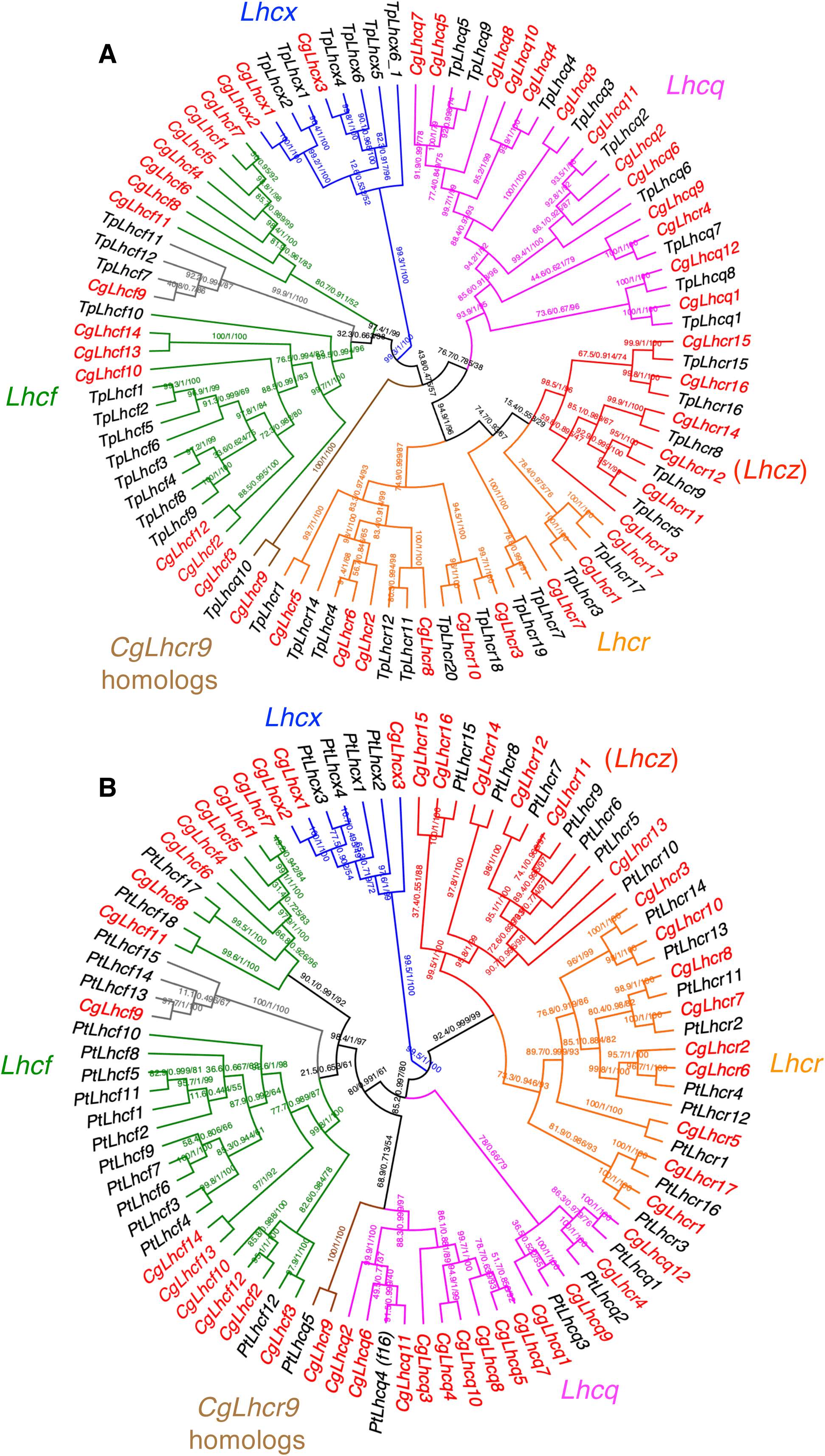
Maximum-likelihood trees of FCPs from *Chaetoceros gracilis* (Cg) and *Thalassiosira pseudonana* (Tp) and from *Chaetoceros gracilis* and *Phaeodactylum tricornutum* (Pt). The trees were inferred using IQ-TREE 2 (Minh et al., 2020). The numbers of supporting values are SH-aLRT support (%)/aBayes support/ultrafast bootstrap support (%). Colors of clades are as follows: magenta, Lhcq subfamily; red, Lhcz subfamily; orange, Lhcr subfamily; brown, CgLhcr9 homologs; green, Lhcf subfamily (CgLhcf9 homolog clade is in gray); blue, Lhcx subfamily. Colors of gene names are as follows: red, *Chaetoceros gracilis* FCP; black, *Thalassiosira pseudonana* FCP. **A**, Maximum-likelihood tree of 46 CgFCPs and 44 TpFCPs. The tree was inferred using the LG+F+R4 model selected with ModelFinder (Kalyaanamoorthy et al., 2017). **B**, Maximum-likelihood tree of 46 CgFCPs and 42 PtFCPs. The tree was inferred using the LG+F+R5 model selected with ModelFinder.

The Lhcr subfamily is a red-algal-type LHC shared among both red algae and red-lineage secondary symbiotic algae. The Lhcr subfamily consists of LHCI in red algae (Pi et al., 2018). The Lhcf subfamily was named after fucoxanthin, while other FCP subfamily proteins also bind it. Lhcf was also named as Fcp; However, this nomenclature is confusing and should be avoided. The Lhcf clade contained a branch of CgLhcf9, in which PtLhcf15 was included as a red-light-induced FCP (**Fig. 2B**). The unique functions of PtLhcf15 under red light conditions have been suggested (Herbstová et al., 2017). Lhcx subfamily proteins are involved in photoprotection through NPQ in diatoms (Buck et al., 2019). This Lhcx subfamily is homologous to Lhcsr, which is also responsible for energy-dependent NPQ (qE) (Tokutsu and Minagawa, 2013; Giovagnetti and Ruban, 2018). The Lhcz subfamily was found in Cryptophyceae, Haptophyta, and Chlorarachniophyta, although its expression, function, and localization are unknown (Koziol et al., 2007). The Lhcz subfamily in diatoms has also been reported by Dittami *et al*. (2010). This Lhcz subfamily was assigned to the Lhcr clade or as a sister clade of the Lhcr subfamily in our phylogenetic trees. Therefore, the systematic names of the Lhcz subfamily are described as Lhcr herein.

The fifth subfamily is Lhcq, a novel FCP subfamily proposed in our previous study (Nagao et al., 2020). The functions of the Lhcq subfamily are unknown. Although Lhcq proteins were not annotated in the model diatoms, Lhcqs were partially annotated as Lhcy proteins by Nymark *et al*. (2013) and Clade V by Hoffman *et al*. (2011). The Lhcq clade was distinguished by high support values (e.g., 93.9/1/95 for SH-aLRT support [%]/aBayes support/ultrafast bootstrap support [%]; **Fig. 2A**). The Lhcq subfamily was more similar to the Lhcf subfamily than to the Lhcr and Lhcx subfamilies based on likelihood mapping analysis (Strimmer and von Haeseler, 1997) (**Supplemental Fig. S1**).

In addition to the five subfamilies, a minor number of FCPs comprised a monophyletic clade containing CgLhcr9. CgLhcr9 homologs also included the protein Pt17531 (protein ID 17531 in the JGI database; *Phaeodactylum tricornutum* CCAP 1055/1 v2.0, https://phycocosm.jgi.doe.gov/Phatr2/Phatr2.home.html). The functions of CgLhcr9 homologs were unknown until their localization in the PSI-FCPI complex was reported (Nagao et al., 2020). CgLhcr9 homologs in *Thalassiosira pseudonana* and *Phaeodactylum tricornutum* were named Lhcq because they were phylogenetically independent of the typical Lhcr subfamily; however, CgLhcr9 itself was still considered a member of the Lhcr subfamily because of its structural composition in PSI-FCPI.

Based on the above phylogenetic analysis, we proposed that 44 TpFCPs and 42 PtFCPs could be renamed into the four subfamily names (Lhcr, Lhcf, Lhcx, and Lhcq; **Supplemental Tables S2, S3**). In particular, TpFCPs and PtFCPs belonging to the Lhcq clade were renamed as Lhcq using our new annotations. Some Lhcrs, previously considered Lhcas (e.g., RefSeq ID: XP_002287377.1 and XP_002289005.1) were renamed as Lhcrs. Consequently, there were nine Lhcrs, 14 Lhcfs, three Lhcxs, six Lhczs, 13 Lhcqs including CgLhcr4, and CgLhcr9 in *Chaetoceros gracilis*; 11 Lhcrs, 12 Lhcfs, six Lhcxs, five Lhczs, nine Lhcqs, and one CgLhcr9 homolog in *Thalassiosira pseudonana*; and nine Lhcrs, 17 Lhcfs, four Lhcxs, seven Lhczs, four Lhcqs, and one CgLhcr9 homolog in *Phaeodactylum tricornutum* (**Fig. 2A, B**).

The centric diatoms *Chaetoceros gracilis* and *Thalassiosira pseudonana* had orthologous gene sets of Lhcr, Lhcz, Lhcq, and CgLhcr9 homologs, whereas some gene duplications and a minor exception, i.e., CgLhcr13 (Lhcz), were absent in *Thalassiosira pseudonana* (**Fig. 2A**). Notably, Lhcf- and Lhcx-type FCPs formed branches within each species, suggesting that Lhcr, Lhcz, Lhcq, and CgLhcr9 homologs may have conserved functions in both species, whereas Lhcf and Lhcx may have been differentiated within each species. A similar tendency was observed between *Chaetoceros gracilis* and *Phaeodactylum tricornutum* (**Fig. 2B**); however, *Phaeodactylum tricornutum* had a smaller number of Lhcq genes compared with *Chaetoceros gracilis* and *Thalassiosira pseudonana*. All PtLhcqs had putative orthologous FCPs in *Chaetoceros gracilis*, although several CgLhcq homologs was missing in *Phaeodactylum tricornutum*. We further extended our phylogenetic analysis to FCPs from other diatoms, such as *Thalassiosira oceanica* (Lommer et al., 2012), *Fistulifera solaris* (Tanaka *et al*., 2015), *Fragilariopsis cylindrus* CCMP1102 (Mock et al., 2017), and *Pseudo-nitzschia multistriata* (**Supplemental Table S5**). The relatively conserved set of Lhcr, Lhcz, and CgLhcr9 homologs was found among all species, except *Pseudo-nitzschia multistriata*, which has only three Lhcr-type FCPs (**Supplemental Fig. S2A–C**), and the completely conserved set of Lhcrs among six species corresponding to those of *Chaetoceros gracilis* is listed in **Supplemental Table S6**. Diverged sets of Lhcf, Lhcq, and Lhcx were observed among other diatoms.

### Localization of five major FCP subfamilies in PSI-FCPI and PSII-FCPII structures of Chaetoceros gracilis

Nagao *et al*. (2020) named FCPI proteins of *Chaetoceros gracilis* based on their localization in the PSI-FCPI structure. Internal FCPs that formed a ring-like structure around the PSI core were named CgLhcr1–10, and peripheral FCPs, which bound the above internal FCPs, were named CgLhcq1–6 (**Fig. 3A**). Among these FCPs, CgLhcr1–3, CgLhcr5–8, and CgLhcr10 belonged to the Lhcr subfamily; CgLhcr4 and CgLhcq1–6 branched into the Lhcq clade; and CgLhcr9 branched into an independent clade (**Fig. 2A, B**). The larger *Chaetoceros gracilis* PSI-FCPI supercomplex reported by Xu *et al*. (2020) also contained CgLhcq9 (FCPI-19), CgLhcq12 (FCPI-2), CgLhcf3 (FCPI-12), another CgLhcq6 (FCPI-20), and CgLhcq5 (FCPI-21; **Fig. 3B**). Moreover, this structure contained three additional FCPs (FCPI-1, -17, and -18); however, the amino acid sequences used for the structural modeling were those from *Phaeodactylum tricornutum* (PtLhcf3/4: XP_002177868.1/XP_002177869.1) and *Fragilariopsis cylindrus* (OEU13194.1, Fracy1:210193 in JGI, and A0A1E7F4Y9 in UniProtKB; OEU18584.1 had a partial sequence of OEU13194.1). OEU13194.1 is a red algal lineage chlorophyll *a*/*b*-binding-like protein (redCAP) and is different from typical LHC proteins, such as FCP (Sturm et al., 2013). Because of the low sequence similarity, redCAP proteins were not obtained in our BLASTP search. If the sequences used for structural modeling were relevant to the actual CgFCP sequences, two FCPs (FCPI-17 and FCPI-18) modeled by PtLhcf3/4 sequences may belong to the Lhcf subfamily. Thus, eight Lhcr-, 11 Lhcq-, one CgLhcr9-, and three Lhcf-type FCPs as well as one unknown FCP may function as light-harvesting antennae for PSI.

**Figure 3.**
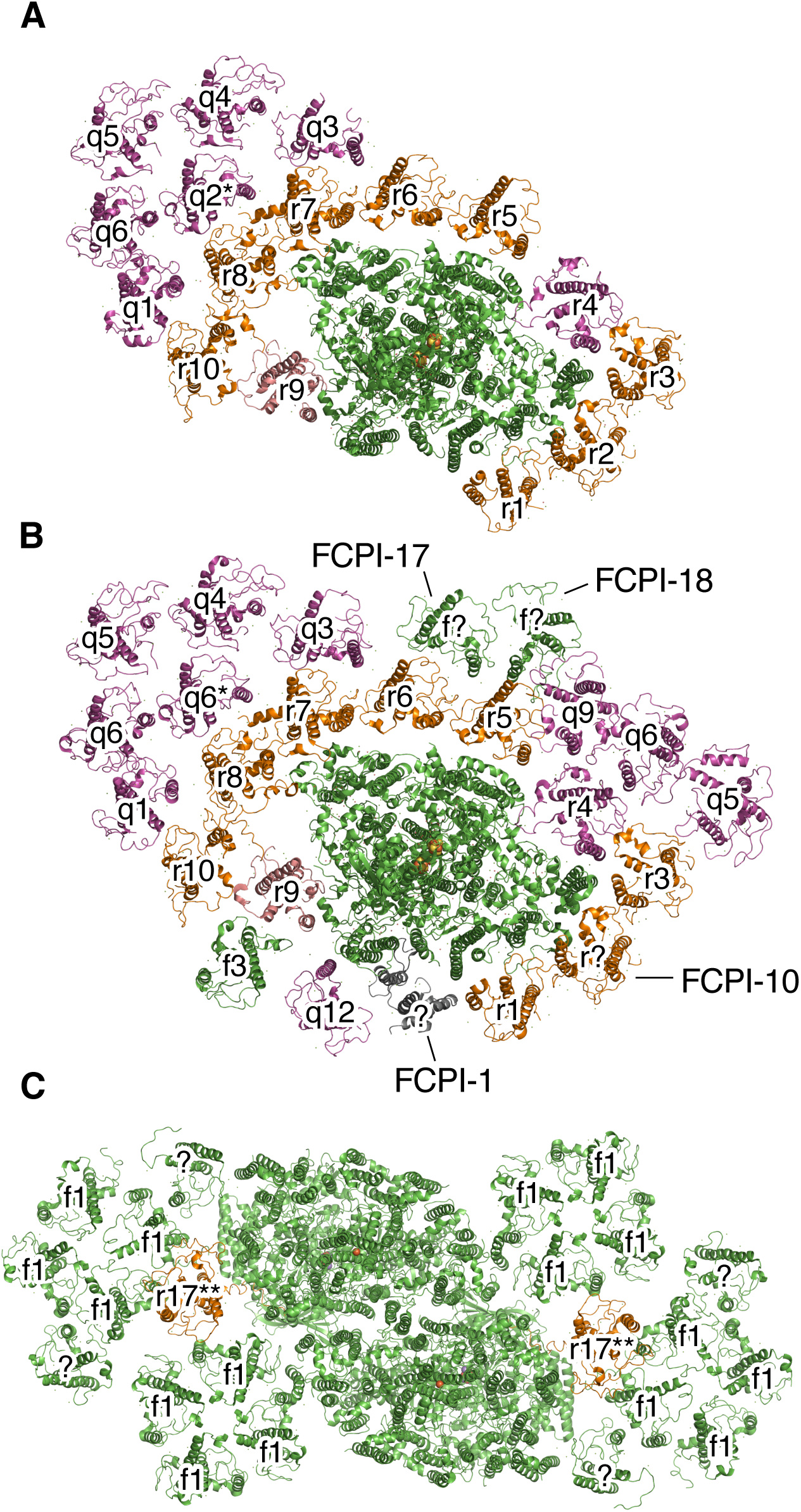
Structural arrangements of the photosystem I-FCPI supercomplex (A, PDB ID: 6L4U; B, PDB ID: 6LY5) and the photosystem II-FCPII supercomplex (C, PDB ID: 6J40) of *Chaetoceros gracilis*. Top view of each supercomplex from the stromal side was depicted using PyMOL (Schrodinger LLC, 2015). The colors of FCPs are indicated as follows: magenta, Lhcq subfamily; red, Lhcz subfamily; orange, Lhcr subfamily; salmon pink, CgLhcr9 homologs; green, Lhcf subfamily. **A**, Sixteen FCPs were assigned in the PSI-FCPI supercomplex. **B**, Twenty FCPs were assigned, among which 24 FCPs were found in the larger PSI-FCPI supercomplex. Four unassigned FCPIs are indicated as Xu *et al*. (2020). CgLhcq2 (q2^*^) was assigned in **A**: 6L4U (Nagao et al., 2020), whereas CgLhcq6 was assigned in **B**: 6LY5 (Xu et al., 2020). **C**, CgLhcf1 tetramers were assigned in the dimeric PSII-FCPII supercomplex (Nagao *et al*., 2019); CgLhcr17 (r17^**^) was assigned in Pi *et al*. (2019). Two FCP monomers in each monomer of the PSII-FCPII were not assigned in both reports. The unassigned FCPs are shown in green.

The positions of the five Lhcrs in red algal PSI-LHCI were similar to those of CgLhcr1, CgLhcr5, CgLhcr6, CgLhcr7, and the unknown FCPI-1 (**Fig. 3B**); however, their orthologous relationships were not supported by our phylogenetic analysis (**Supplementary Fig. S3A, B**). Interestingly, CgLhcr9 bound to the PSI core in an orientation opposite that of other endogenous FCPI proteins. The unique binding mode of CgLhcr9 in PSI-FCPI is related to its separation from other medial FCPI proteins in the phylogenetic tree (**Fig. 2A, B**). The position occupied by the Lhcq protein in the PSI-FCPI of *Chaetoceros gracilis* is completely absent in the PSI-LHCI of the red alga *Cyanidioschyzon merolae* (Nagao *et al*., 2020; Pi *et al*., 2018). Therefore, the Lhcq subfamily, including Lhcr4, is likely a new addition from secondary endosymbiosis.

*Chaetoceros gracilis* PSII-FCPII formed dimers and had two tetramers and three monomers of FCPs per PSII core (Nagao *et al*., 2019; Pi *et al*., 2019) (**Fig.3C**). Nagao *et al*. (2019) revealed that two tetramers consisted of CgLhcf1, and Pi *et al*. (2019) reported that the center monomer of three monomers was Lhca2, which was renamed CgLhcr17 based on the systematic nomenclature in this study. The presence of the Lhcr-type FCP in the PSII-FCPII complex suggests its special function in light energy transfer. Because of the resolution limit, the molecular identities of the other two FCP monomers in the PSII-FCPII complex are still unknown.

### Putative organization of FCPs surrounding photosystems in other diatoms

The detailed structures of photosystems from other diatoms have not been elucidated. However, an orthologous set of Lhcr-type FCPs, including CgLhcr4 from the Lhcq subfamily, except for CgLhcr17 homologs, was detected in purified PSI complexes from both centric *Thalassiosira pseudonana* and pennate *Phaeodactylum tricornutum* using mass spectrometry (Lepetit *et al*., 2010; Grouneva *et al*., 2011; Ikeda *et al*., 2013; Calvaruso *et al*., 2020) (**Figs. 4, 5**). Notably, the PSI-FCPI complex of *Thalassiosira pseudonana* reported by Calvaruso *et al*. (2020) lacks TpLhcr18 and TpLhcr20, corresponding to CgLhcr3 and CgLhcr10, respectively (**Fig. 4**). These FCPs would be detached during the isolation process. TpLhcr17, an ortholog of CgLhcr17, was detected in a PSII-FCPII fraction (Calvaruso *et al*., 2020). Thus, most Lhcrs have specific and conserved functions as antennae for PSI, with the exception of CgLhcr17 homologs for PSII.

**Figure 4.**
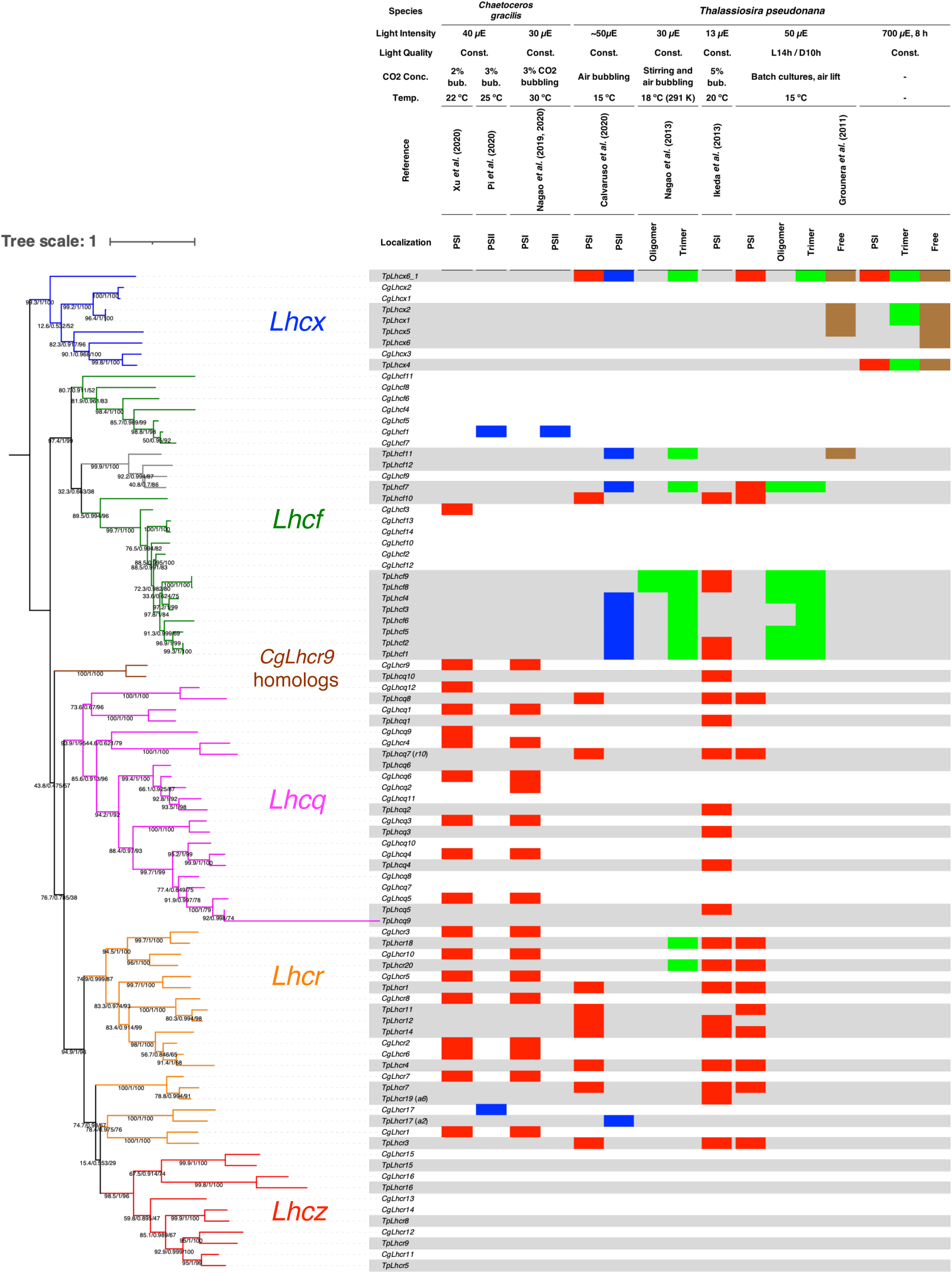
Maximum-likelihood tree of FCPs from *Chaetoceros gracilis* (Cg) and *Thalassiosira pseudonana* (Tp) combined with the table showing their previous detection in purified protein complexes. The trees were inferred using IQ-TREE 2 (Minh et al., 2020) with the LG+F+R4 model selected using ModelFinder (Kalyaanamoorthy et al., 2017). Numbers of supporting values are SH-aLRT support (%)/aBayes support/ultrafast bootstrap support (%). The tree was rerooted with the Lhcx subfamily. Detection of FCPs in each fraction or band is indicated by colored boxes as follows: red, PSI; blue, PSII; green, trimer; brown, free. Colors of clades are as follows: magenta, Lhcq subfamily; red, Lhcz subfamily; orange, Lhcr subfamily; brown, CgLhcr9 homologs; green, Lhcf subfamily (CgLhcf9 homolog clade is in gray); blue, Lhcx subfamily.

**Figure 5.**
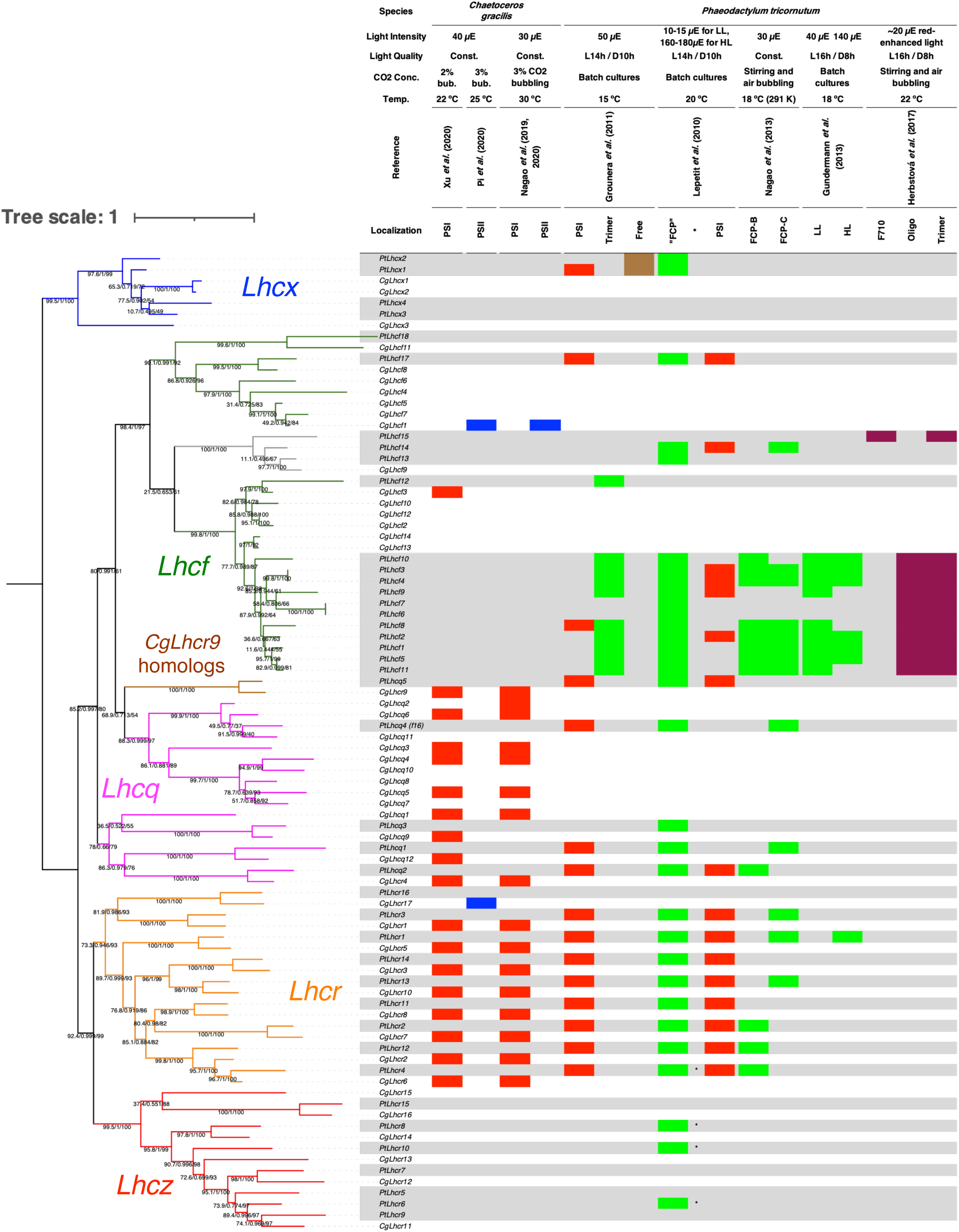
Maximum-likelihood tree of FCPs from *Chaetoceros gracilis* (Cg) and *Phaeodactylum tricornutum* (Pt) combined with the table showing their previous detection in purified protein complexes. The trees were inferred using IQ-TREE 2 (Minh et al., 2020) with the LG+F+R5 model selected using ModelFinder (Kalyaanamoorthy et al., 2017). Numbers of supporting values are SH-aLRT support (%)/aBayes support/ultrafast bootstrap support (%). The tree was rerooted with the Lhcx subfamily. Detection of FCPs in each fraction or band is indicated by colored boxes as follows: red, PSI; blue, PSII; green, trimer; brown, free; purple, FCPs induced by red light. *PtLhcr4, PtLhcr6, PtLhcr8, and PtLhcr10 proteins could be detected with a few peptides under HL, while they were completely missing under LL. Colors of clades are as follows: magenta, Lhcq subfamily; red, Lhcz subfamily; orange, Lhcr subfamily; brown, CgLhcr9 homologs; green, Lhcf subfamily (CgLhcf9 homolog clade is in gray); blue, Lhcx subfamily.

In the centric diatom *Thalassiosira pseudonana*, almost full sets of Lhcq subfamily proteins, except for TpLhcq9 and TpLhcq6, were detected in the PSI-FCPI band separated by native PAGE (Ikeda *et al*., 2013), whereas only TpLhcq7 and TpLhcq8, corresponding to CgLhcr4 and CgLhcq12 located in the inner-ring PSI-FCPI, were detected in two other studies (Grouneva *et al*., 2011; Calvaruso *et al*., 2020) (**Fig. 4**). In the pennate diatom *Phaeodactylum tricornutum* (**Fig. 5**), only PtLhcq2, orthologous to CgLhcr4, was detected (Lepetit *et al*., 2010; Grouneva *et al*., 2011), suggesting a conserved role of CgLhcr4 homologs in PSI-FCPI. By contrast, PtLhcq1 and PtLhcq4, corresponding to CgLhcq12 and CgLhcq10, respectively, were detected in PSI-FCPI in one study (Grouneva *et al*., 2011). The discrepancy between the two studies may be related to differences in cultivation conditions or purification processes. TpLhcq10 and PtLhcq5—CgLhcr9 homologs—were detected in both *Thalassiosira pseudonana* and *Phaeodactylum tricornutum* PSI-FCPI (Lepetit *et al*., 2010; Grouneva *et al*., 2011; Ikeda *et al*., 2013). Taken together, the organization of FCPs in the inner-ring FCPI surrounding PSI was similar among *Chaetoceros gracilis, Thalassiosira pseudonana*, and *Phaeodactylum tricornutum*, whereas the composition of outwardly bound FCPs was diverse among diatom species.

CgLhcf1 forms homotetramers and serves as the main antennae in the PSII-FCPII of *Chaetoceros gracilis* (Nagao et al., 2020). In *Thalassiosira pseudonana* (**Fig. 4**), TpLhcf1-7 and TpLhcf11 were detected in the PSII-FCPII fraction (Calvaruso et al., 2020), consistent with the function of Lhcf-type FCPs as the main antennae for PSII. Additionally, TpLhcf1-7 and TpLhcf11 were also detected as “FCP trimers” in other studies (Grouneva *et al*., 2011; Nagao *et al*., 2013), indicating that FCPII consisting of Lhcfs was loosely attached to PSII and easily detached during isolation. In *Phaeodactylum tricornutum* (**Fig. 5**), isolation of the PSII-FCPII complex has not been reported, although free “FCP trimers” have been reported in several studies (Lepetit *et al*., 2010; Grouneva *et al*., 2011; Gundermann *et al*., 2013; Nagao *et al*., 2013), potentially representing detachment of FCPII from PSII. Notably, the exact oligomeric state of the freely isolated “FCP trimer” is unknown, although the structures of FCP tetramers (Nagao *et al*., 2019; Pi *et al*., 2019) and dimers (Wang et al., 2019) have been elucidated, and the trimeric form has been observed in cryo-EM single particle analysis (Arshad et al., 2021).

As described previously, CgLhcf3 was putatively assigned to *Chaetoceros gracilis* PSI-FCPI (Xu et al., 2020). TpLhcf10 was detected in the *Thalassiosira pseudonana* PSI-FCPI fraction (Calvaruso et al., 2020), but not in PSII-FCPII nor the “trimer” (Grouneva *et al*., 2011; Nagao *et al*., 2013; Calvaruso *et al*., 2020). Therefore, some Lhcf-type proteins may serve as FCPI in both species; nevertheless, the diversification of Lhcf-type seems to have occurred independently (**Figs. 2A, 4**). In *Phaeodactylum tricornutum*, PtLhcf2, PtLhcf3/4, PtLhcf9, PtLhcf14, and PtLhcf17 were detected in the PSI-FCPI fraction (Lepetit *et al*., 2010), whereas PtLhcf8 and PtLhcf17 were detected elsewhere (Grouneva *et al*., 2011) (**Fig. 5**). These Lhcf-type FCPs may compensate for the smaller number of Lhcqs in PSI-FCPI of *Phaeodactylum tricornutum*. Further structural analysis of PSI-FCPI of *Phaeodactylum tricornutum* is required.

### Motif analysis of FCPs

LHC/FCP is a pigment-protein complex with three transmembrane alpha helices (α1, α2, and α3) (Engelken et al., 2010) or helices B, C, and A (Kühlbrandt et al., 1994; Bassi et al., 1999) from the N-terminus. In all FCPs, the α1 and α3 helices have sequence similarity and highly conserved glutamate (E64 and E163 in CgLhcf1) and arginine (R69 and R168 in CgLhcf1) residues that interact in an interhelix manner (E64-R168 and E163-R69); thus, the interaction between the α1 and α3 helices seems to be stabilized, as indicated in green plant LHCII (Engelken et al., 2010) (**Fig.6**). The highly conserved glutamates of the α1 and α3 helices also coordinate Chls in the conserved composition.

**Figure 6.**
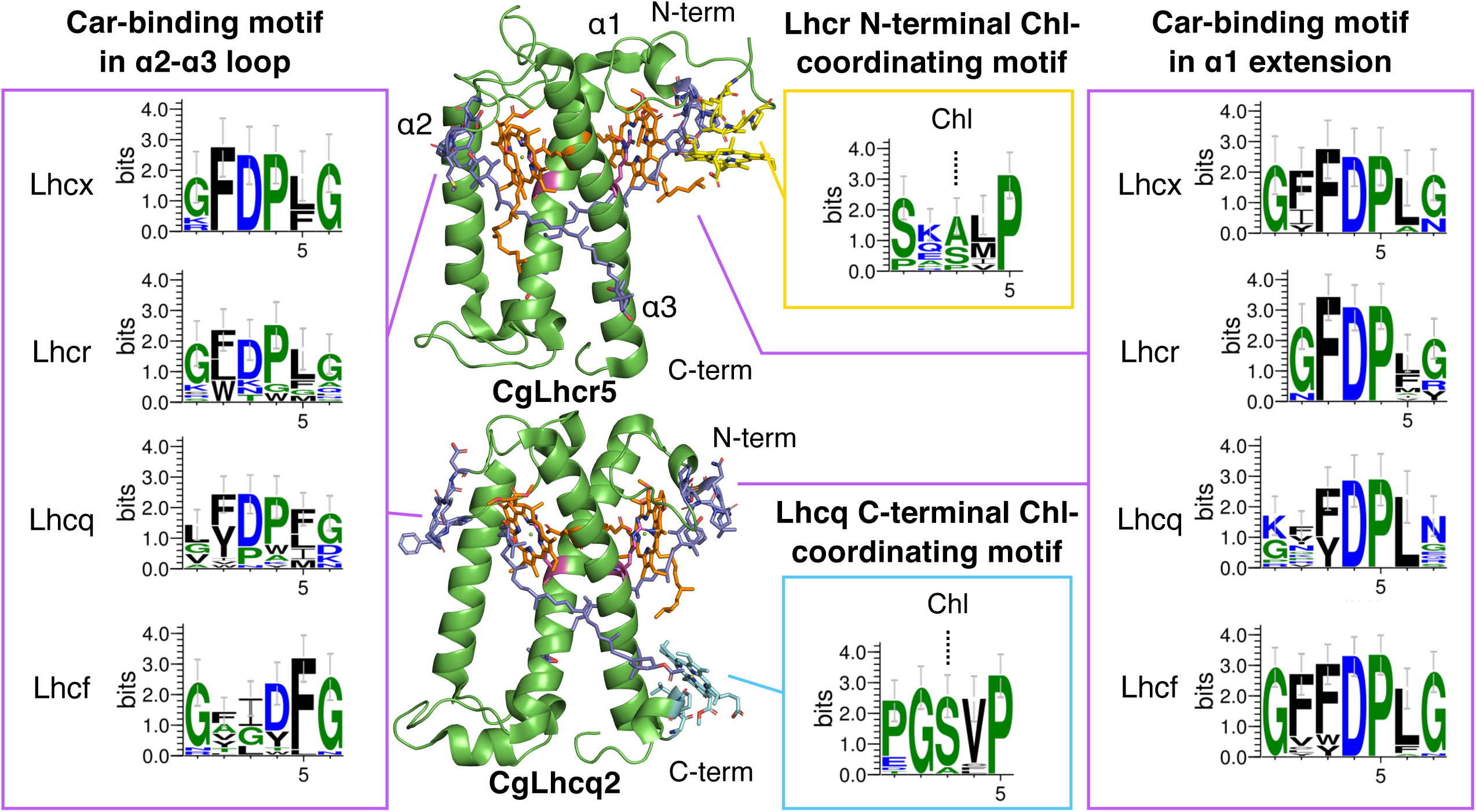
Structural localization and sequence logos of the pigment-binding motifs in FCPs. CgLhcr5 and CgLhcq2 structures from *Chaetoceros gracilis* PSI-FCPI (PDB ID: 6L4U) were depicted using PyMOL (Schrodinger LLC, 2015). The cartoon model shows the side view of each FCP with the stromal side up. Not all Chls or carotenoids are shown. The amino acid residues and their coordinating or binding pigments are shown as stick models: carotenoid-binding motifs and carotenoids, purple; glutamate, orange; arginine, magenta; Lhcr N-terminal Chl-coordinating motif SX[S/A]X[L/M]P, yellow; Lhcq C-terminal Chl-coordinating motif PGSVP, cyan. Motif logos were created using WebLogo 3.7.4 (Crooks et al., 2004).

For further motif analysis, multiple expectation maximization for motif elicitation (MEME, version 5.3.0) (Bailey et al., 2009) was performed using translated sequences of FCP genes from *Chaetoceros gracilis* and *Thalassiosira pseudonana* (**Supplemental Fig. S4**). LHC/FCP had two common carotenoid (Car)-binding motifs in the extended sequence of the N-terminal side of the α1 helix (α1 extension) and in the loop region between helices α2 and α3 (α2–α3 loop), with GFDPLG or similar sequences in some varieties (**Fig. 6**) (Bassi *et al*., 1999; Engelken, Brinkmann, and Adamska 2010). These conserved Car-binding motifs were detected by MEME as motif-2, -5, or -9 in the α1 extension and in the α2–α3 loop of Lhcr, Lhcx, Lhcz, and Lhcq subfamilies (**Supplemental Figs. S4, S5**). The typical GFDPLG sequence was conserved in the α1 extension of Lhcr and in the α2–α3 loop of Lhcr and Lhcx (**Fig. 6**). Although the motif in the α2–α3 loop of Lhcr was less conserved, only Lhcr, the most ancestral FCP subfamily in diatoms, had a typical Car-binding sequence in both regions.

The Car-binding motif in the α1 extension of Lhcf and Lhcx contained GFFDPLG, with addition of phenylalanine to the typical sequence. The corresponding region of Lhcq (14/22 Lhcq sequences from *Chaetoceros gracilis* and *Thalassiosira pseudonana*) was [K/G]X[F/Y]DPLN, which had the same number of amino acids as Lhcf and Lhcx. By contrast, CgLhcq9, CgLhcq12, CgLhcr4, and its homolog TpLhcq7 had various deletions or insertions before conserved aspartic acid residues in the Car-binding motif, and CgLhcq1, CgLhcq3, and their homologs had different residues between [K/G] and X (**Supplemental Fig. S6A**). The motif in the α2–α3 loop of Lhcq showed minor variations with X[F/Y]DP[F/L]G, whereas that of Lhcf was largely different from the typical Car-binding motif represented by GXXDFG, lacking proline and having different locations of aspartic acid residues. Thus, Lhcf was identified as the newest subfamily among the five major FCP subfamilies.

The conserved Chl-coordinating motif of SX[S/A][L/M]P, a part of MEME motif-6 (**Supplemental Fig. S4**) was located on the stromal side (**Fig. 6**) and contained only the N-terminal sequence of Lhcr proteins, except for those of CgLhcr3 and CgLhcr10. Interestingly, FCPs of Lhcf, Lhcx, Lhcz, and Lhcq subfamilies, except for CgLhcx3 and TpLhcx4–6, completely lacked this motif in their N-terminal region, indicating that this motif was lost during the functional differentiation of FCPs in diatoms. In the Lhcr subfamily, this motif coordinates Mg^2+^ of Chl *a* via the carbonyl oxygen in the peptide bond of S/A at 2.1–2.6 Å distance (Nagao *et al*., 2020; Xu *et al*., 2020). These Chls are located between CgLhcr1/2, -2/3, -5/6, -6/7, -7/8, and -8/10 and may contribute to the assembly and stabilization of FCPI and to energy transfer to adjacent Chls. CgLhcr10 had a PEPIP sequence instead of SX[S/A][L/M]P, and its glutamate residue coordinated Chl *a* at 2.4 Å via its side chain. The N-terminal region coordinating Mg^2+^ of Chl *a* may be an ancestral property of red algal LHC because this motif was also observed in Lhcr1 of *Cyanidioschyzon merolae* (Pi et al., 2018). Similarly, another sequence motif coordinated Mg^2+^ of Chl *a* in the LHC proteins of green-lineage plants: Lhcas (PDB ID: 6JO5), CP26, and CP29 (PDB ID: 6KAD) had PX[W/F]LP in *Chlamydomonas reinhardtii* (Sheng et al., 2019; Su et al., 2019; Suga et al., 2019), and LhcbMs (PDB ID: 6KAD) had [P/A][K/L][F/W]LGP (Sheng et al., 2019). Each motif in the LHC proteins of *Chlamydomonas reinhardtii* coordinated Mg^2+^ of Chl *a* via its carbonyl oxygen in the main chain of W/F at a 2.1–2.8 Å distance.

CgLhcr3 and CgLhcr10 are unique among the diatom Lhcr subfamily because of differences in the N-terminal Chl-coordinating motif and their positions in the genome. Both CgLhcr3 and CgLhcr10 were encoded in scaffold00008 in the *Chaetoceros gracilis* draft genome, with their 5′ ends placed head-to-head. Homologous genes for *CgLhcr3* and *CgLhcr10* (*TpLhcr18* and *TpLhcr20*, EJK71515.1 and RJK71517.1, and *PtLhcr14* and *PtLhcr13* in *Thalassiosira pseudonana, Thalassiosira oceanica*, and *Phaeodactylum tricornutum*, respectively) were also arranged in a head-to-head manner, suggesting differentiation from other Lhcr subfamily genes at an early stage of diatom diversification.

The C-terminal conserved motif PGSVP, a part of MEME motif-11 (**Supplemental Fig. S6B, C**), on the lumenal side of the Lhcq subfamily coordinates Mg^2+^ of Chl *a* via the carbonyl oxygen of the peptide bond from the serine residue (Nagao *et al*., 2020) (**Fig. 6**). This PGSVP motif is conserved in the Lhcq subfamilies of *Chaetoceros gracilis, Thalassiosira pseudonana*, and *Phaeodactylum tricornutum*. The MEME motif-11, including this PGSVP motif, was also assigned to some Lhcf subfamily proteins (**Supplemental Fig. S6B**). However, the region of Lhcf proteins did not contain the first proline. Chl 316 with adjacent Chls and carotenoids in Lhcq proteins is involved in the excitation energy transfer between FCPs toward the circumferential direction in PSI-FCPI (Nagao et al., 2020). The PGSVP motif corresponds to the TGKGP motif in LHCs of *Chlamydomonas reinhardtii* in multiple sequence alignment; however, the latter motif does not coordinate Mg^2+^ of Chl *a*. Therefore, the PGSVP motif specific to Lhcq was also obtained during FCP differentiation to retain additional Chl.

## Discussion

### Distributions and functions of Lhcrs

The Lhcr subfamily, which includes LHCI in red algae, is present in a wide variety of red algal lineages and contains an independent clade of the Lhcz subfamily (**Fig. 7A, Supplemental Figs. S3A, B, S7A– I**). However, LHCI of the two red algae analyzed herein did not show orthologous relationships with the Lhcrs of secondary endosymbiotic algae in the red algal lineage. By contrast, the secondary symbiotic algae in the red algal lineage had gene sets similar to those of Lhcrs with phylogenetic relevance to those in diatoms, suggesting that functional differentiation of the Lhcr subfamily occurred during secondary symbiotic events in the red algal lineage.

**Figure 7.**
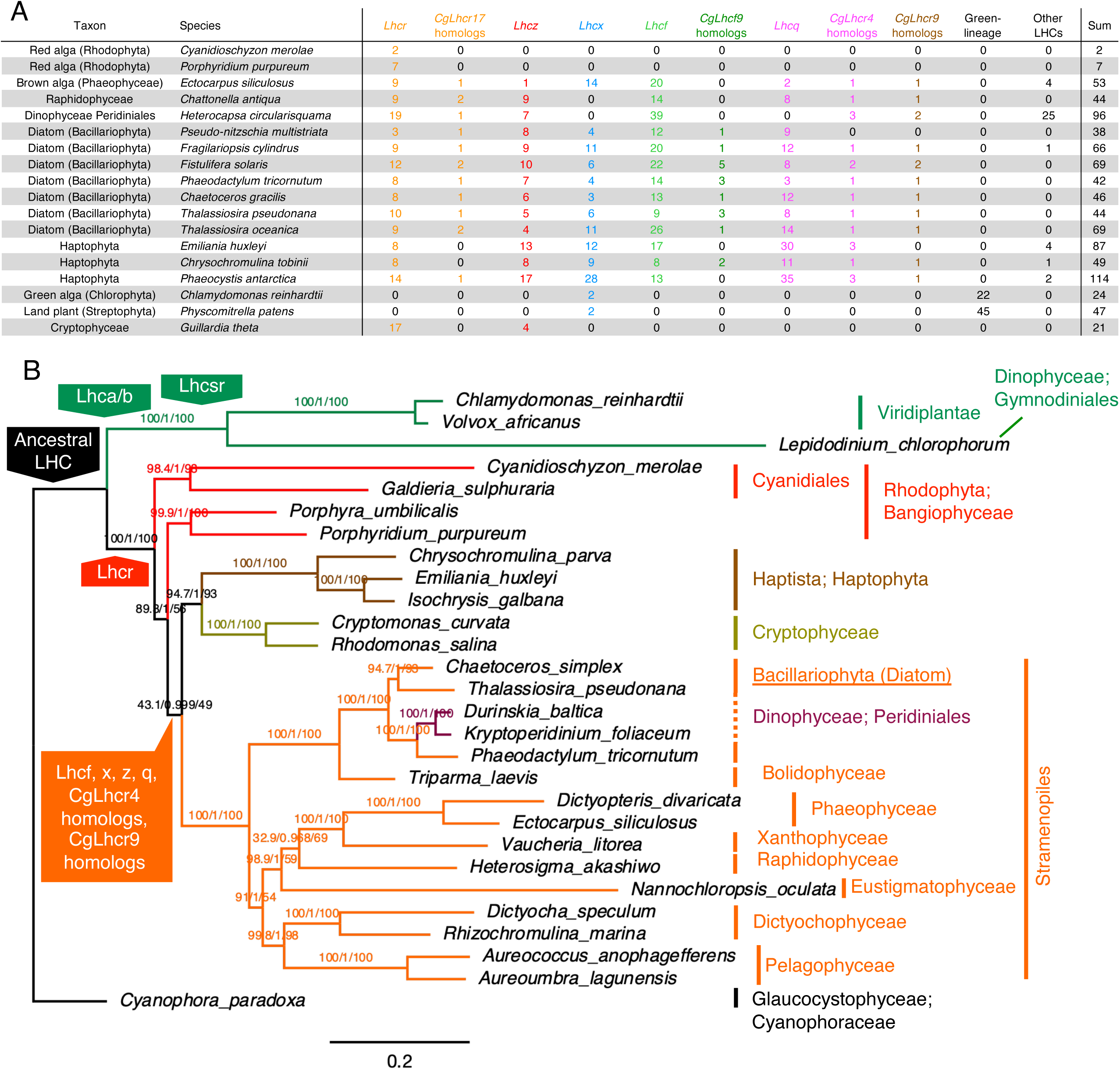
Distribution of LHC/FCP subfamilies among red-lineage algae and hypothesis of these acquisitions based on the phylogenetic tree of chloroplast genes. **A**, Numbers of FCP/LHC belonging to each subfamily detected from each species. **B**, Maximum-likelihood tree generated using chloroplast-encoded genes from various algal species indicating the estimated acquisition point of each LHC/FCP subfamily. The tree was constructed using models selected with ModelFinder (Kalyaanamoorthy et al., 2017) for each gene. The tree was rerooted with Graucocystophyceae. Numbers of supporting values are SH-aLRT support (%)/aBayes support/ultrafast bootstrap support (%).

This Lhcr differentiation may have occurred gradually during red algae evolution; most diatoms shared an orthologous gene set of Lhcrs (**Supplemental Table S6**), although homologs of CgLhcr2 and CgLhcr6 were not clearly distinguished in the phylogenetic trees. Other Stramenopiles have homologs of CgLhcr10 in Phaeophyceae and Raphidophyceae; however, they do not have homologs of CgLhcr3, and those of CgLhcr2 and CgLhcr6 were not clearly separated in their trees. In Haptophyta, the presence or number of homologs for CgLhcr2/CgLhcr6, CgLhcr3/CgLhcr10, and CgLhcr7/CgLhcr8 differed between species; *Phaeocystis antarctica* lacked CgLhcr2/CgLhcr6, CgLhcr3/CgLhcr10, and CgLhcr7/CgLhcr8, whereas *Emiliania huxleyi* lacked CgLhcr3/CgLhcr10 and *Chrysochromulina tobinii* had homologs for CgLhcr2/CgLhcr6, but only one gene for CgLhcr7/CgLhcr8 and CgLhcr3/CgLhcr10. These differences could be related to incompleteness of genome or transcriptome analyses and may cause structural differences in the inner ring of FCPI surrounding PSI among the secondary symbiotic algae in the red algal lineage.

Interestingly, one central monomer (Nagao *et al*., 2019; Pi *et al*., 2019) of the three monomers in diatom PSII-FCPII was identified as CgLhcr17, the only FCPII belonging to the Lhcr subfamily. CgLhcr17 homologs were conserved in Stramenopiles, including *Pseudo-nitzschia multistriata*, Phaeophyceae, and Raphidophyceae, and only *Phaeocystis antarctica* from Haptophyta had a CgLhcr17 homolog (**Fig. 7A, Supplemental Fig. S7**). In the Haptophyta analyzed in this study, *Chrysochromulina tobinii* and CgLhcr17 homologs were missing, and in the other Haptophyta *Emiliania huxleyi*, the phylogeny among CgLhcr1, CgLhcr17, and XP_005769864.1 was unclear. CgLhcr17 homologs belong to clade II described by Hoffman *et al*. (2011), which includes several haptophyte Lhcrs. Therefore, photosynthetic Stramenopiles and some Haptophytes may have an Lhcr-type FCP for PSII-FCPII. The specific function of CgLhcr17 in PSII-FCPII is unknown.

CgLhcr9 was designated Lhcr based on its binding location in *Chaetoceros gracilis* PSI-FCPI (Nagao et al., 2020) and belongs to the independent clade from Lhcr and other subfamilies in the phylogenetic tree. Clade VI described by Hoffman *et al*. (2011) corresponded to CgLhcr9 homologs, which were conserved in diatoms (**Fig. 7A, Supplemental Table S6**), other Stramenopiles, and two Haptophytes; however, the other Haptophyta *Emiliania huxleyi* showed unclear phylogeny around CgLhcr9.

### Diversification of the Lhcf subfamily within each species

The Lhcf subfamily is a group of FCPs that accumulates abundantly and includes FCPs assigned to the diatom PSII-FCPII. Free or “trimeric” fractions of FCPs are mainly composed of Lhcf-type FCPs, and few Lhcfs may also attach to the larger diatom PSI-FCPI (Xu et al., 2020). Phylogenetic analyses suggested that Lhcf subfamily proteins were diversified within each species, indicating that changes in the Lhcf subfamily may be essential for adapting to light environments. One of the subordinate groups of Lhcf subfamilies is a group that includes CgLhcf9 and PtLhcf15, which is independent from other subclades of Lhcf subfamilies. PtLhcf15 constitutes red-shifted FCPs induced by red light exposure (Herbstová et al., 2017). In this clade, *Chaetoceros gracilis* had only CgLhcf9, which may be induced by red light. CgLhcf9 homologs were identified in both diatoms and Haptophytes (**Figs. 2A, 2B, 7A, Supplemental Fig.S2A–D**), indicating that the Lhcf subfamily is also involved in chromic adaptation. However, CgLhcf9 homologs were not detected in the Dinophyceae Peridiniales transcriptome.

### Lhcx subfamily for energy-dependent NPQ

The Lhcx subfamily is responsible for energy-dependent NPQ (qE) components in diatoms and other red-lineage secondary symbiotic algae (Giovagnetti and Ruban, 2018) and is widely conserved among secondary symbiotic red-lineage algae. In green algae and moss, the homologous subfamily is called Lhcsr or LI818 protein (Zhu and Green, 2008), which is also responsible for qE (Bailleul et al., 2010). Lhcx and Lhcsr subfamilies are classified into Clade V by Hoffman *et al*. (2011). Vascular plants lack Lhcsr subfamily proteins but have PsbS, which belongs to the LHC superfamily as an NPQ entity (Li *et al*., 2000). Generally, Lhcx subfamily proteins are upregulated with increasing light intensity at the mRNA level, contributing to light intensity-dependent induction of NPQ, whereas only *PtLhcx1* is expressed constitutively under dark conditions (Bailleul et al., 2010). Calvaruso *et al*. (2020) identified TpLhcx6_1 in both PSI and PSII fractions of thylakoid membranes separated in the centric diatom *Thalassiosira pseudonana* under both low and high light conditions, TpLhcx4 in PSI from samples treated with high light, and TpLhcx1/2 in free fractions from samples treated with high light. Grouneva *et al*. (2011) identified TpLhcx4 in PSI from samples treated with high light, TpLhcx1/2 in free fractions from samples treated with high light, and PtLhcx1 in PSI and PtLhcx2 in free fractions of thylakoid membranes from *Phaeodactylum tricornutum*.

Orange carotenoid-binding protein (OCP) is responsible for NPQ in cyanobacteria and is activated under high light conditions and connects to phycobilisome, membrane anchored light-harvesting pigment-protein complexes (Joshua *et al*., 2005; Thurotte *et al*., 2015; Kirilovsky, 2007; Kirilovsky and Kerfeld, 2016). Primary symbiotic red-lineage algae have NPQ capacity (Schubert et al., 2011; Wu, 2016; Álvarez-Gómez et al., 2019), whereas both OCP homologs and Lhcx/Lhcsr subfamilies are absent in red algal genomes (Tanaka et al., 2004; Bhattacharya et al., 2013). Thus, the molecular entity of NPQ in red algae remains unknown.

### The Lhcq subfamily

The Lhcq subfamily is a new subfamily of FCPs comprising the outer belt of FCPs in the PSI-FCPI complex of *Chaetoceros gracilis* (Nagao et al., 2020; Xu et al., 2020). *Chaetoceros gracilis* and *Phaeodactylum tricornutum* have different features of excitation energy transfer from FCP to PSI with different amounts of low-energy Chls in PSI and/or FCPI (Nagao et al., 2018; Nagao et al., 2019b; Tanabe et al., 2020); this may be related to the reduced number of Lhcq proteins in *Phaeodactylum tricornutum* compared with *Chaetoceros gracilis*.

Among Lhcqs in *Chaetoceros gracilis*, CgLhcr4 was considered Lhcr because of its location in the inner ring of FCPs in PSI-FCPI, interacting with PsaB, PsaF, and Psa28. Unlike other Lhcq subfamily proteins, homologs of CgLhcr4 were widely conserved in the secondary symbiotic algae of the red lineage, i.e., Stramenopiles and Haptophytes (**Fig. 7A**). Therefore, the FCP compositions in the inner-ring PSI-FCPI should be conserved in Stramenopiles and Haptophytes. Photosynthetic Stramenopiles other than diatoms also had Lhcq subfamily proteins in addition to homologs of CgLhcr4; however, these are not orthologous to diatom Lhcqs. Haptophyta had Lhcq homologs belonging to a large sister clade of CgLhcq4, CgLhcq5, CgLhcq7, CgLhcq8, and CgLhcq10, in addition to CgLhcr4 homologs (**Supplemental Fig. S7C, E, H**). This suggested that peripheral region PSI-FCPI supercomplexes of other Stramenopiles and Haptophyta may be different from those of diatoms, indicating that CgLhcr4 homologs may be the oldest members of the Lhcq subfamily.

### Hypothesis of diversification of FCP/LHC subfamilies in the red lineage

Secondary symbiotic algae in the red algal lineage have obtained genes targeted for or encoded in the plastids from the symbiont of an ancient red alga. Red algae harvest light mainly via Lhcrs as antennae for PSI and use phycobilisome as antennae for PSII. However, phycobilisome genes are absent in the genomes of all secondary symbiotic algae in the red algal lineage. In diatoms, PSI uses Lhcrs, Lhcqs, the CgLhcr9 homolog, and several Lhcfs, whereas PSII uses the CgLhcr17 homolog and Lhcfs.

The diversification of LHC/FCP subfamilies was coupled with the phylogenetic diversification of red algal lineages. However, their phylogeny is complicated by symbiotic gene transfer (SGT) via primary, secondary, and tertiary endosymbiosis and horizontal gene transfer (HGT) (Keeling, 2013). Indeed, Dorrell *et al*. (2017) reported that 25% of plastid-targeted genes of red-lineage secondary symbiotic algae were derived from the green lineage. Thus, we constructed a phylogenetic tree of 65 single-copy orthologous genes encoded in plastid genomes (**Supplemental Table S7**) detected by OrthoFinder (**Fig. 7**). In this tree, secondary symbiotic algae in the red algal lineage were suggested as monophyletic groups, consistent with a report by Kim *et al*. (2017). Haptophyta was inferred to be a sister group of Stramenopiles, consistent with the phylogenetic tree of the nuclear genome (Burki *et al*., 2016). The Dinophyceae Peridiniales was located in a clade of diatoms, as suggested by Horiguchi and Takano (2006). Overall, our phylogenetic tree using plastid-encoded genes did not contradict previously presented trees.

Confusion regarding the phylogenic relationships of FCP/LHC subfamilies has hindered our understanding of the diversification process of red-lineage FCP/LHC subfamilies. Our FCP/LHC phylogenetic analysis including *Chlamydomonas reinhardtii* and *Physcomitrella patens* from green-lineage plants revealed that all subfamilies from the red lineage, except for Lhcx (Lhcsr) subfamilies, were independent from green-lineage LHCs, such as Lhca and Lhcb subfamilies, with high support values. Our analysis also suggested that the Lhcr subfamily was the most relevant subfamily and that Lhcq and Lhcf subfamilies were not derived from green-lineage LHC genes through HGT or SGT. By contrast, the process of acquiring Lhcxs was not clarified.

Similarities between Lhcq and Lhcf subfamilies were supported by likelihood mapping using 46 CgFCPs and 44 TpFCPs, suggesting that Lhcr, Lhcf, Lhcx, and Lhcq subfamilies could be grouped as (Lhcr, Lhcx)-(Lhcq, Lhcf). These findings were also supported by similarities in the pigment-binding motifs of Lhcqs and Lhcfs; the N-terminal motif in the stromal side coordinating Chl *a* was conserved in Lhcr, Lhca, and Lhcb subfamilies but absent in both Lhcq and Lhcf subfamilies, whereas the C-terminal motif in the lumenal side coordinating Chl *a* was conserved in Lhcq and some Lhcf proteins. There were also differences between Lhcq and Lhcf subfamilies; for example, the Car-binding motif in the α2–α3 loop of Lhcq had proline, similar to other subfamilies, whereas that of Lhcf subfamily did not have proline. Thus, Lhcq and Lhcf may have a common ancestor derived from an ancestral Lhcr subfamily protein, and the Lhcf subfamily may have been derived from the Lhcq subfamily.

Based on these findings, we propose the following process for LHC/FCP diversification. First, the common ancestor of secondary symbiotic algae (excluding Cryptophyceae) acquired Lhcr subfamily genes from the red algal symbiont and diversified not only Lhcr genes but also CgLhcr9 homologs and Lhcq genes, including CgLhcr4 homologs. This diversification enlarged the antenna size of the PSI. During this process, CgLhcr17 was derived from one of the Lhcr genes to fit into the PSII core instead of phycobilisomes, and Lhcf subfamily proteins diverged from the Lhcq subfamily, generating several monomers and tetramers and attaching to PSII as a major light-harvesting antenna. This hypothesis was supported by analyses of LHC/FCP distributions and their functions in various secondary symbiotic algae in the red lineage, particularly in those other than diatoms, using many genome and transcriptome sequencing results from various species.

## Conclusion

Our draft genome and transcriptome analyses suggested that *Chaetoceros gracilis* had 46 FCP genes, classified into five major subfamilies, i.e., Lhcr, Lhcf, Lhcx, Lhcz, and Lhcq, and one minor subfamily, i.e., CgLhcr9. FCPs of the inner light-harvesting ring of the PSI-FCPI complex were composed of Lhcrs, including CgLhcr9 and several Lhcqs, and were highly conserved in other diatom species. By contrast, Lhcfs, some of which were found in the PSII-FCPII complex, seemed to be diversified in each diatom species, and the number of Lhcqs differed among species. This indicated that diversification of Lhcf and Lhcq contributed to species-specific adaptations to the light environment. Other algae in Stramenopiles and Haptophyta possess the five major subfamilies and CgLhcr9 homologs. Therefore, FCP/LHC diversification would have occurred in the common ancestral origin of red lineage algae.

## Materials and Methods

### Diatom cultivation

The marine centric diatom *Chaetoceros gracilis* (UTEX LB 2658) was used for all analyses. Cell cultures were prepared in f/2 artificial seawater (Guillard, 1975) under 30 µmol photons m^-2^ s^-1^ at 20°C with continuous shaking at 100 rpm. Additional cell culture for IsoSeq analysis was performed in artificial seawater under 30 µmol photons m^-2^ s^-1^ at 30°C with continuous bubbling of air containing 3% (v/v) CO_2_ (Nagao *et al*., 2007).

### Genome sequencing and draft genome assembly

Genomic DNA was isolated as previously described (Fischer et al., 1999) and analyzed using the Genome Sequencer FLX+ System (GS FLX+; Roche Diagnostics, Basel, Switzerland), Genome Analyzer GAIIx (Illumina, Inc., San Diego, CA, USA), and Hiseq (Illumina, Inc.).

The sequencing library for GS FLX+ was prepared using the Paired End Library Preparation Method Manual (20 kb and 8 kb Span). The obtained library was amplified by emulsion polymerase chain reaction (PCR) using a GS FLK Titanium SV/LV emPCR Kit (Lib-L; Roche Diagnostics), added to a GS FLK Titanium PicoTiterPlate (Roche Diagnostics), and sequenced using the GS FLX+ System with a GS FLK Titanium Sequencing Kit XL+. The sequences of 744,262 reads containing 319,847,738 bases (454 BaseCaller 2.6 in GS FLX+ system software) were used for the assembly process.

GAIIx sequencing was based on TruSeq DNA Sample Preparation v2 Guide Rev. A using a TruSeq DNA Sample Preparation v2 Kit. Sequences were called based on Genome Analyzer IIx User Guide version A and TruSeq SBS Kit v5 Reagent Preparation Guide (for Genome Analyzer) version C. Sequences were processed following the Consensus Assessment of Sequence and Variation (CASAVA) v1.8 User Guide version B; 22,251,716 reads were processed.

The DNA sequences used in Hiseq were prepared using TruSeq. In total, 26,854,816 reads of 101 paired bases were obtained. Adapter sequences were eliminated using Cutadapt v1.1 (Martin, 2011). Low-quality bases were trimmed using Trimmomatic v0.32 (Bolger, Lohse, and Usadel, 2014); paired reads of every 5 bases with a higher average quality score of 30 and whose lengths were longer than 74 survived; 16,222,538 reads survived (average length: 100.1).

Genome assembly was performed with all sequence data obtained from GS FLX+, GAIIx, and Hiseq using GS *De Novo* Assembler version 2.8 (Roche Diagnostics) with the following assembly parameters: -nrm/-het/-a0/-ml80%/-mi90/-urt/-large.

### RNA extraction

RNA extraction was performed using a RNeasy kit (Qiagen Inc., Valencia, CA, USA) with some modifications. Cells were centrifuged, and pellets were suspended in 600 µL RLT buffer containing 1% (v/v) β-mercaptoethanol. The suspension was sonicated 10 times for 0.2–0.3 s each time using Handy Sonic UP-21P (Tomy, Japan) and centrifuged for 3 min at 15,000 rpm at room temperature. The supernatant was transferred to a new 1.5-mL microtube and mixed with the same volume of 70% ethanol. The mixture was further processed using the standard RNeasy protocol.

### RNA sequencing

The library for RNA sequences was prepared using a Directional mRNA-Seq Library Prep. Pre-Release Protocol Rev.A with TruSeq RNA Sample Prep Kit and TruSeq Small RNA Sample Prep Kit (Illumina Inc.). The library was reverse transcribed, amplified by PCR with primers containing indexes, and purified by 6% agarose gel electrophoresis. Clustering was performed using cBot User Guide version F and TruSeq SR Cluster Kit v2 Reagent Preparation Guide (for cBot) version C and analyzed with Genome Analyzer IIx User Guide version A and TruSeq SBS Kit v5 Reagent Preparation Guide (for Genome Analyzer) version C. Base calling and processing were performed based on CASAVA v1.8 version B. The sequences were single reads of 75 bases. In total, 110,067,972 reads were obtained. RNA sequences were mapped to the genome using Hisat2 (Kim *et al*., 2019).

### Gene prediction

Genes in the *Chaetoceros gracilis* draft genome were predicted using BRAKER2 (Hoff et al., 2016) with AUGUSTUS trained with RNA-seq mapping data. FCP genes were manually curated using RNA-seq mapping. Gene prediction of the chloroplast genome was performed using DOGMA (https://dogma.ccbb.utexas.edu/). The information for predicted genes is available at ChaetoBase (https://chaetoceros.nibb.ac.jp/).

### Iso-Seq

The Iso-seq libraries from two samples prepared under different cultivation conditions were generated according to the protocol provided by Pacific Biosciences (PN 101-763-800 Version 1; CA, USA), using NEBNext Single Cell/Low Input cDNA Synthesis & Amplification Module (New England Biolabs, MA, USA), Iso-Seq Express Oligo Kit, SMRTbell Express Template Prep Kit 2.0, and Barcoded Overhang Adapter Kit. The 5′ and 3′ primers GCAATGAAGTCGCAGGGTTGGG and AAGCAGTGGTATCAACGCAGAGTAC were used. Each Iso-Seq library was sequenced using Pacific Bioscience Sequel II. From each library, 1,968,854 and 1,035,268 raw reads were obtained, with average lengths of 2,744 and 3,310 bases, respectively.

The raw sequences were processed to ccs reads using SMRT Link v8.0.0, according to the SMRT Link User Guide (v8.0) version 09. The ccs reads were refined, and FLNC reads were created using IsoSeq3 refine (Gordon et al., 2015). FLNC reads were then clustered using IsoSeq3 (Gordon et al., 2015) with the verbose option and “use qvs”. The refined FLNC reads were also clustered using isONclust v0.0.6 (Sahlin and Medvedev, 2019). Open reading frames were extracted using TransDecoder v5.5.0 (Haas et al., 2013) from respective clustered reads processed by IsoSeq3 or isONclust, and two sets of translated sequences were obtained from each of two sets of raw reads.

### Genome and transcriptome quality assessment

Basic statistics were analyzed using Seqkit (Shen et al., 2016). To assess genome or transcriptome assembly and gene prediction completeness, BUSCO (v4.0.6) (Seppey *et al*., 2019) was performed in protein mode. The BUSCO lineage datasets used in our analyses were selected based on their taxonomy. Stramenopile_odb10 was selected for diatoms, Raphidophyceae, and Pheaophyceae (brown alga).

### Acquisition of the FCP sequences of Chaetoceros gracilis and model diatoms

Forty-four sequences of CgFCPs were collected from the draft genome data using BLASTP 2.10.0 similarity search (Altschul et al., 1990). In each BLASTP search, 30 and 39 known TpFCPs and PtFCPs collected from RefSeq were used as queries, and the expectation value (E-value) threshold was set to 1e-05. BLASTP searches were also conducted for each set of IsoSeq translated sequences generated by IsoSeq3 or isONclust. *CgLhcf13* (GenBank ID: LC647435) and *CgLhcf14* (GenBank ID: LC647436) were identified from the Iso-Seq sequences. Using 46 CgFCPs, known TpFCPs, and PtFCPs as queries, BLASTP similarity searches were performed to obtain FCP sequences from *Thalassiosira pseudonana* and *Phaeodactylum tricornutum* genomes (Armbrust *et al*., 2004; Bowler *et al*., 2008) with an E-value threshold of 1e-5. The 44 TpFCPs and 42 PtFCPs were phylogenetically analyzed for classification of their genes into different subfamilies, revised based on their phylogeny.

The lists of gene names (**Supplemental Table S3, S4**) were created based on UniProtKB (https://www.uniprot.org/help/uniprotkb).

### Acquisition of FCP sequences from other diatoms and Haptophytes and LHC sequences from red algae

The sets of translated sequences of other diatoms were obtained from the genome assemblies of *Thalassiosira oceanica* (Lommer et al., 2012), *Fistulifera solaris* (Tanaka et al. 2015), *Fragilariopsis cylindrus* CCMP1102 (Mock et al., 2017), and *Pseudo-nitzschia multistriata*. Sets of translated sequences of other red-lineage microalgae were obtained from the following genome assemblies: Phaeophyceae, *Ectocarpus siliculosus* (Cock et al., 2010); three Haptophyta; *Emiliania huxleyi* (Read et al., 2013), *Phaeocystis antarctica* CCMP1374, and *Chrysochromulina tobinii* (Hovde et al., 2015); Cryptophyceae, *Guillardia theta* CCMP2712 (Curtis et al., 2012); and two Rhodophyta, red alga *Porphyridium purpureum* (Bhattacharya et al., 2013) and *Cyanidioschyzon merolae* (Tanaka *et al*., 2004). Sets of translated sequences derived from the RNA-seq data of Raphidophyceae *Chattonella antiqua* and Dinophyceae Peridiniales *Heterocapsa circularisquama* were obtained from the database for research in harmful algal blooms (Shikata *et al*., 2019). Genome assemblies of green-lineage organisms have also been used, including *Chlamydomonas reinhardtii* (Merchant et al., 2007) and *Physcomitrella patens* (Rensing et al., 2008). The accession numbers and URLs are summarized in **Supplemental Table S5**. BLASTP 2.10.0 similarity searches with 1e-5 as the E-value threshold, using 46 CgFCPs, 44 TpFCPs, and 42 PtFCPs as a query set, were conducted for the translated sequences of each genome. The identical LHC/FCP sequences were removed using CD-HIT v4.8.1 (Fu *et al*., 2012) with an identity threshold of 1.0.

### Maximum likelihood phylogenetic analysis

Multiple sequence alignments were constructed using MAFFT-LINSI v7.4 (Katoh and Standley, 2013). Alignments were trimmed using ClipKIT (Steenwyk et al., 2020) with the “kpic-gappy” method, and maximum likelihood phylogenetic trees were constructed using IQ-TREE2 v2.0.7 (Minh et al., 2020). Ultrafast bootstrap (UFBoot2) (Hoang et al., 2018) approximation based on the model selected by ModelFinder (Kalyaanamoorthy et al., 2017) and the SH-like approximate likelihood ratio test (Guindon et al., 2010) were performed with 1000 replications in IQ-TREE2. aBayes test (Anisimova et al., 2011) was also performed. All trees were rerooted with Lhcx clade and drawn using FigTree (v1.4.4, http://tree.bio.ed.ac.uk/software/figtree/) or iTOL (5.7, https://itol.embl.de/) (Letunic and Bork, 2019). Likelihood mapping of Lhcq, Lhcx, Lhcf, and Lhcr (excluding Lhcz) subfamilies was performed using 46 CgFCPs and 44 TpFCPs. CgLhcr9 homologs and Lhcz subfamily sequences were ignored in this analysis.

### Motif analysis of Chaetoceros gracilis FCPs

MEME (v5.3.0) (Bailey *et al*., 2009) was performed with translated sequences of 46 CgFCP and 44 TpFCPs to search 20 motifs. In MEME, the distribution of motifs was not limited, motif lengths were limited from 6 to 50, and other parameters were set to default. Alignment of *Chaetoceros gracilis* and *Thalassiosira pseudonana* FCPs, also used in the phylogenetic analysis, was applied to generate the amino acid sequence logos of the Car-binding motifs and Chl-coordinating motifs in Lhcr, Lhcf, Lhcx, and Lhcq. The logos were visualized using WebLogo 3.7.4 (Crooks et al., 2004). The logo of the Car-binding motifs in the N-terminal extension of the α1 helices (α1 extensions) of Lhcr, Lhcf, Lhcx, and Lhcq were generated without using CgLhcr3, CgLhcr4, CgLhcf11, TpLhcx5, CgLhcq1, CgLhcq3, CgLhcq9, CgLhcq12, TpLhcq1, TpLhcq3, TpLhcq7, TpLhcq9, and TpLhcr18. As a result, 14 out of 22 Lhcqs were used to generate this logo. The logo of the Car-binding motif in the loop region between helices α2 and α3 (α2–α3 loop) was generated without using the CgLhcf9 homolog clade, CgLhcf3, CgLhcf4, TpLhcr4, TpLhcr7, TpLhcr14, TpLhcf6, TpLhcf10, TpLhcx6_1, and TpLhcq10 because of sequence divergence. The logo of the Chl-coordinating motif in the N-terminal extension of Lhcr was generated using the alignment of the proximal Lhcr subfamily, excluding CgLhcr3 and its homolog TpLhcr18. The logo of the Chl-coordinating motif in the C-terminal sequence of Lhcq was generated with alignment of the Lhcq subfamily, excluding TpLhcq9.

### Visualization of the PSI-FCPI and PSII-FCPII structures

PSI-FCPI, PSII-FCPII supercomplexes and FCP structures were visualized using the PyMOL Molecular Graphics System (Version 2.3.0 or 2.4.0 Schrödinger, LLC, https://pymol.org/2/).

### Phylogenetic analysis of chloroplast genomes

The set of translated sequences from chloroplast genomes was used for species phylogenetic analysis. The genomes were selected from NCBI (https://www.ncbi.nlm.nih.gov/) RefSeq or GenBank. All chloroplast genomes used in each analysis are listed in **Supplemental Table S5**, with corresponding accession numbers. Single-copy orthologs among each chloroplast genome were extracted using OrthoFinder v2.5.1 (Emms and Kelly, 2019). Every single-copy ortholog set was aligned using MAFFT v7.4 with the auto option and then trimmed using TrimAl (Capella-Gutiérrez *et al*., 2009) with the automated1 option. The chloroplast phylogenetic tree was inferred using IQ-TREE2 with models automatically selected for each partition of the trimmed alignments. Bootstrap resampling was performed internally using UFBoot with 1000 replicates. Each tree was drawn using FigTree software. Glaucocystophyceae *Cyanophora paradoxa* was used as an outgroup in the chloroplast tree.

## Accession Numbers

Sequence data from this article can be found in our in-house database (https://chaetoceros.nibb.ac.jp/) or in the DDBJ Sequence Read Archive (DRA) under accession numbers DRA012660 (genome sequencing), DRA012661 (RNA-Seq), and DRA012662 (Iso-Seq).

## Supplemental Data

**Supplemental Figure S1**. Likelihood mapping of Lhcr, Lhcq, Lhcf, and Lhcx subfamilies (Strimmer and von Haeseler, 1997).

**Supplemental Figure S2**. Maximum-likelihood trees of FCPs/LHCs from *Chaetoceros gracilis* and other diatoms.

**Supplemental Figure S3**. Maximum likelihood tree of FCPs/LHCs from *Chaetoceros gracilis* and red algae.

**Supplemental Figure S4**. Maximum-likelihood tree of FCPs from *Chaetoceros gracilis* and *Thalassiosira pseudonana* showing the localization of the motifs generated by MEME.

**Supplemental Figure S5**. MEME motif logos contained the conserved carotenoid-binding motif “GFDPLG” with adjacent MEME motif logos.

**Supplemental Figure S6**. Specific motifs of Lhcq subfamily proteins: varieties of carotenoid-binding motifs and the novel C-terminal chlorophyll binding motif “PGSVP”.

**Supplemental Figure S7**. Maximum-likelihood phylogenetic tree of FCPs/LHCs from *Chaetoceros gracilis* and red- and green-lineage species.

**Supplemental Table S1**. List of assemblies used in FCP/LHC detection with BUSCO scores and lineages.

**Supplemental Table S2**. List of *Chaetoceros gracilis* FCPs with gene IDs or accession IDs.

**Supplemental Table S3**. List of all FCPs from *Thalassiosira pseudonana* with revised gene names.

**Supplemental Table S4**. List of all FCPs from *Phaeodactylum tricornutum* with revised gene names.

**Supplemental Table S5**. List of RefSeq or GenBank accession IDs or other references used to obtain the FCP/LHC sequences.

**Supplemental Table S6**. The conserved FCP set of diatoms, including the FCPs assigned to *Chaetoceros gracilis* photosystems.

**Supplemental Table S7**. List of RefSeq or GenBank accession IDs used to infer phylogenetic tree of chloroplast genes.

## Acknowledgements

Computational resources were provided by the Data Integration and Analysis Facility, National Institute for Basic Biology. We would like to thank Editage (www.editage.com) for English language editing.

## Competing interests

The authors declare no competing interests.

## Notes

**Funding information** This work was supported in part by the JST ALCA (grant nos. JPMJAL1105, JPMJAL1608 [K. I. and Y. K.]), by the JSPS KAKENHI (grant nos. JP20H031160 [K.I.], JP20K06528, JP21K19085, JP20H02914 [R.N.], and JP17H06433 [J.R.S.]), and by a Collaborative Research Program from National Institute for Basic Biology (grant no. 21-306 [K.I., Y.K., and I.U.]).

https://chaetoceros.nibb.ac.jp/

### Competing Interest Statement

The authors have declared no competing interest.

