## Supplemental Figures & Tables for "Molecular phylogeny of fucoxanthin-chlorophyll *a/c* proteins from *Chaetoceros gracilis* and Lhcq/Lhcf diversity"

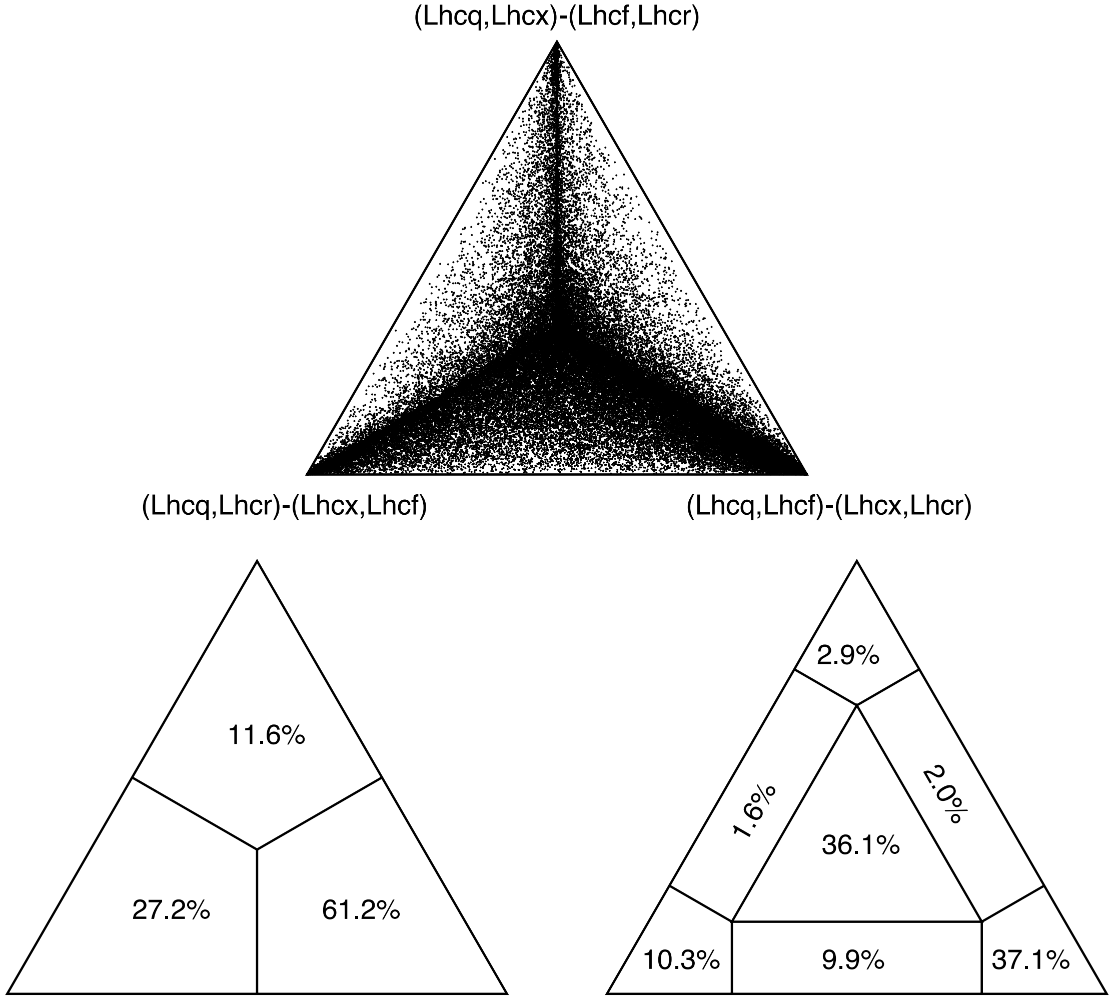


**Supplemental Figure S1. Likelihood mapping of Lhcr, Lhcq, Lhcf, and Lhcx subfamilies (Strimmer and von Haeseler, 1997).** The mapping was inferred using IQ-TREE 2 (Minh *et al.*, 2020) based on the alignment of *Chaetoceros gracilis* and *Thalassiosira pseudonana* FCPs using the VT+F+R4 model selected with ModelFinder (Kalyaanamoorthy *et al.*, 2017).

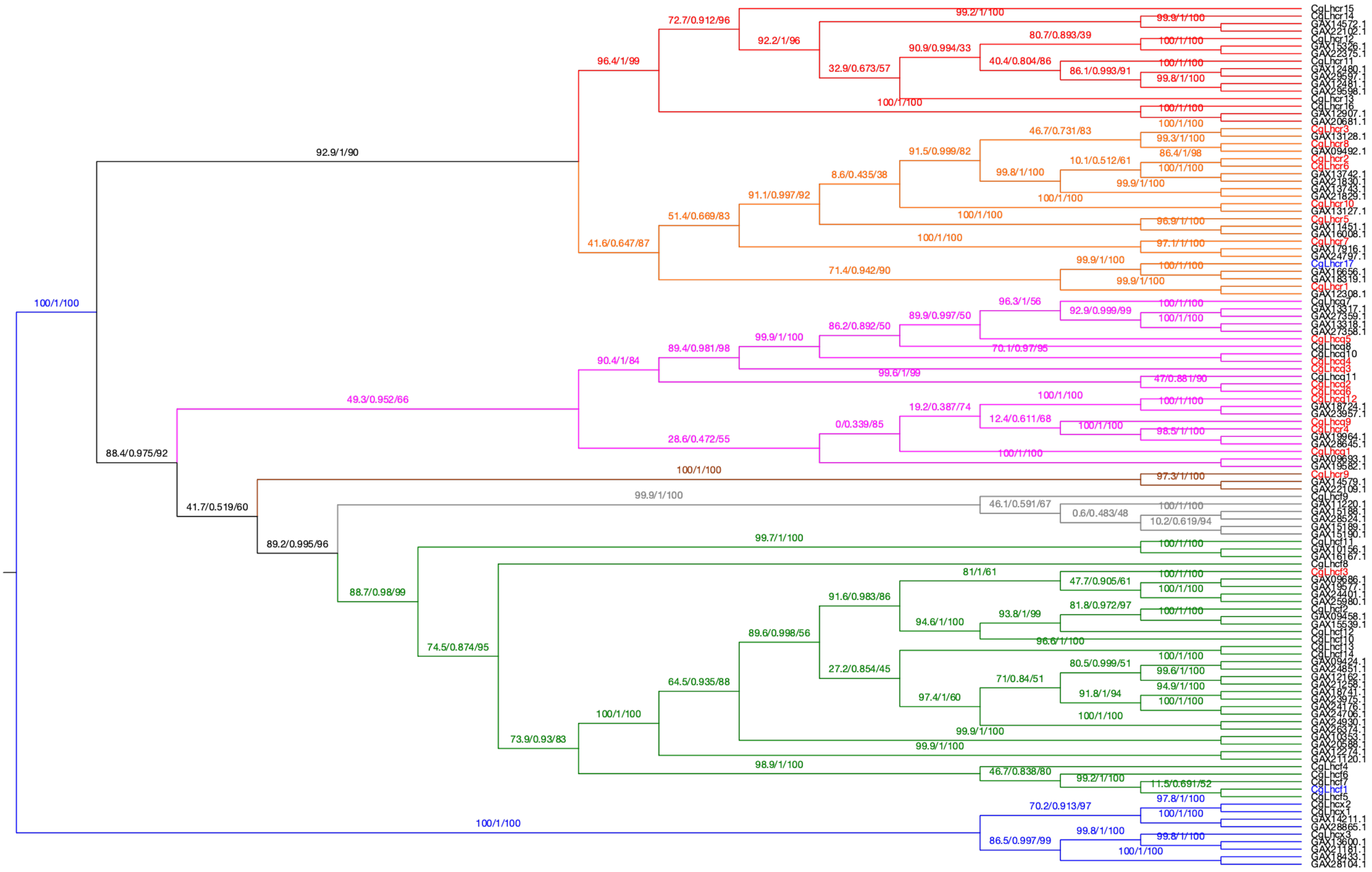

*Fistulifera solaris*

Supplemental Figure S2A

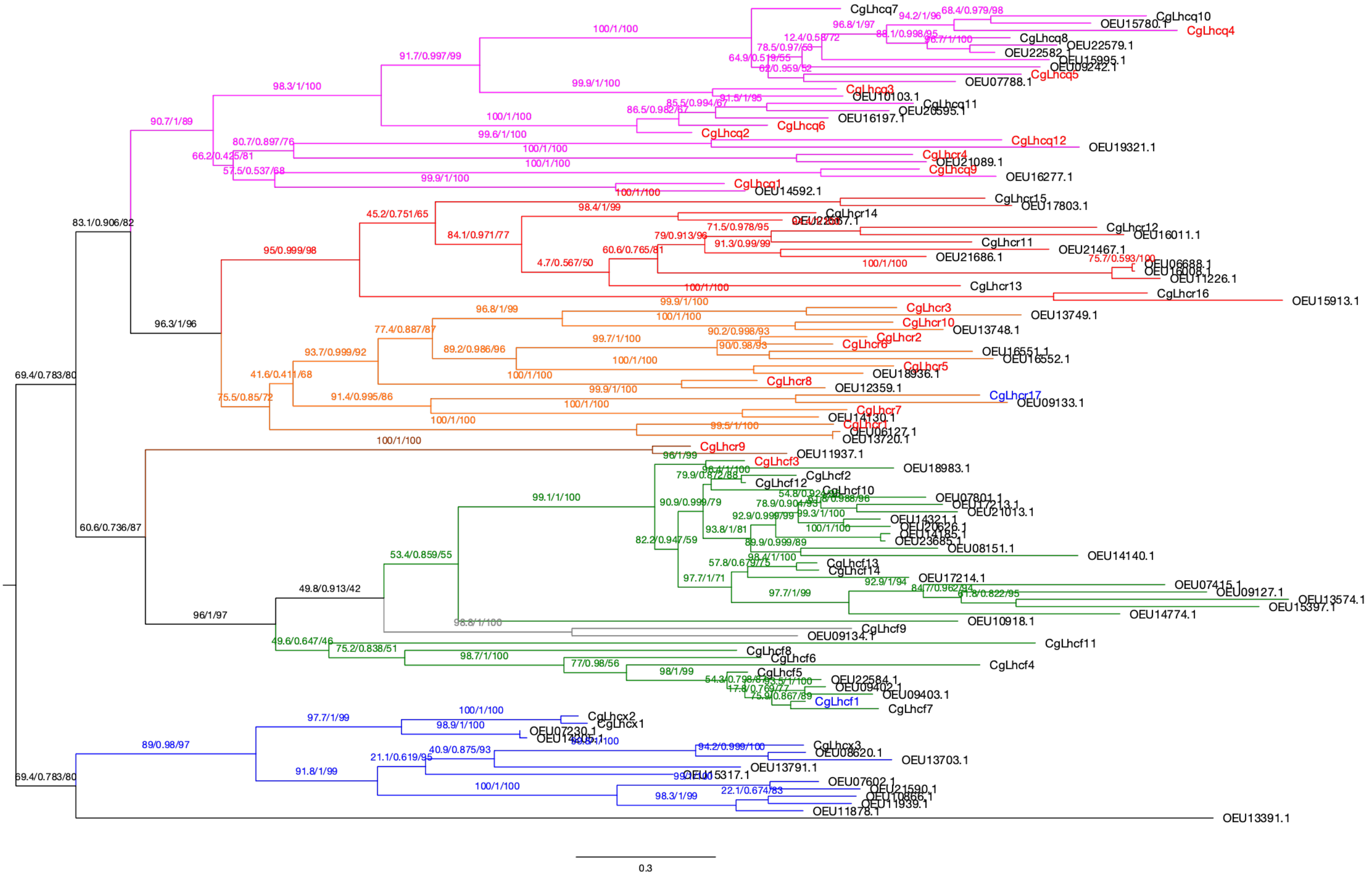

*Fragilariopsis cylindrus*

Supplemental Figure S2B

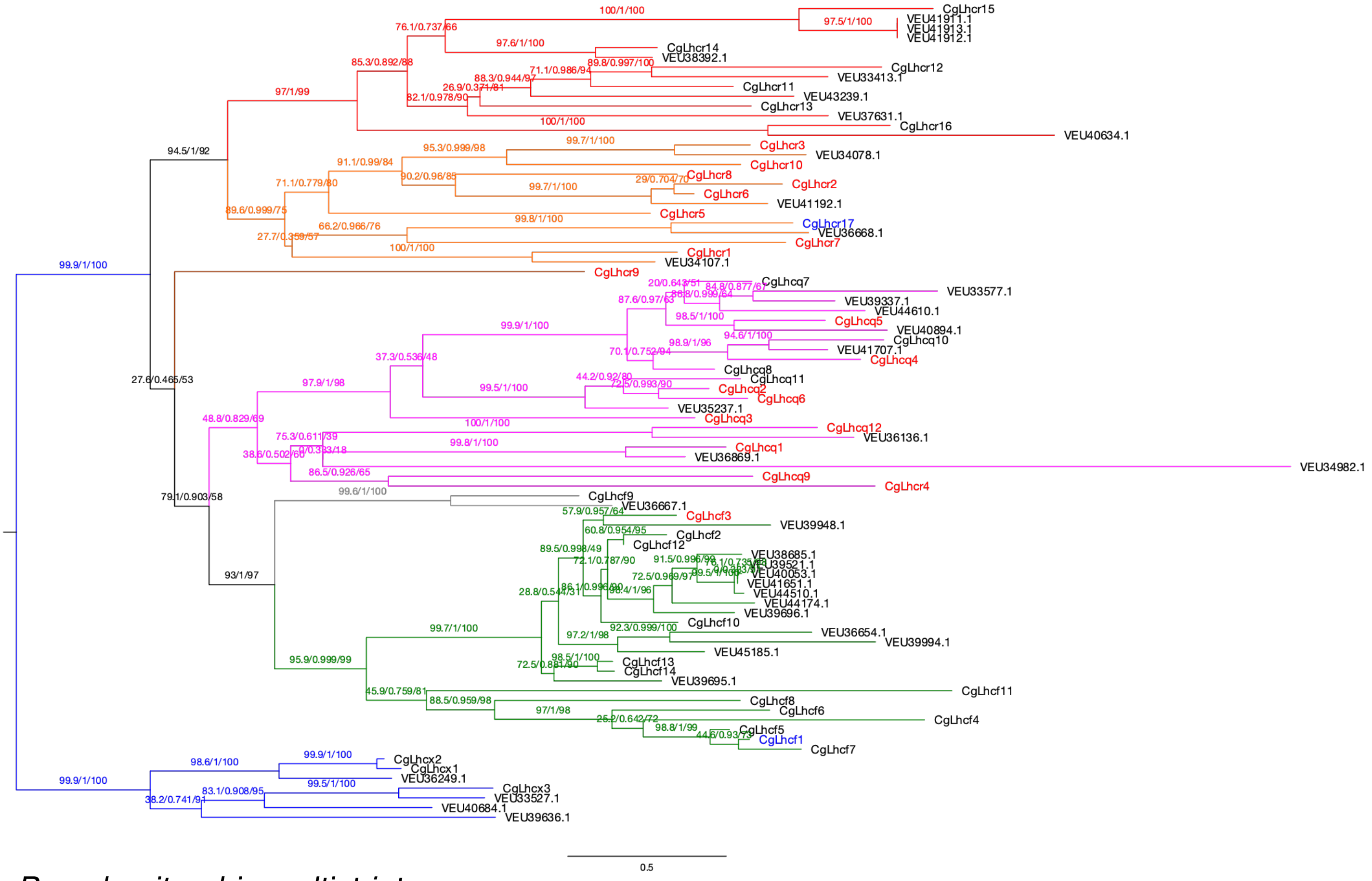


*Pseudo-nitzschia multistriata*

Supplemental Figure S2C

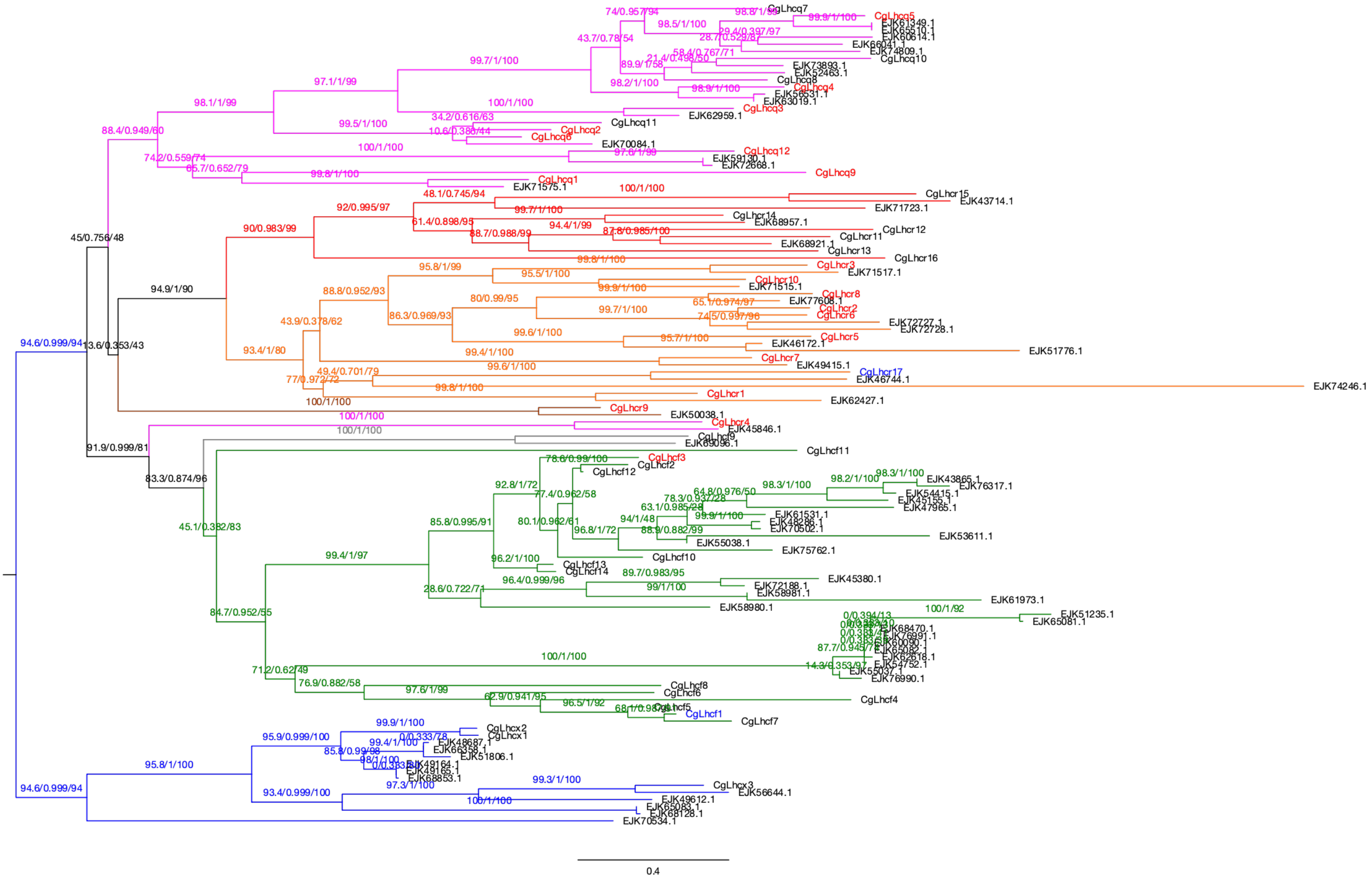


*Thalassiosira oceanica*

Supplemental Figure S2D

**Supplemental Figure S2. Maximum-likelihood trees of FCPs/LHCs from *Chaetoceros gracilis* and other diatoms.** The trees were inferred using IQ-TREE 2 (Minh *et al.*, 2020). Each gene is labeled with GenBank accession ID. The numbers of supporting values are SH-aLRT support (%)/aBayes support/ultrafast bootstrap support (%). Colors of clades are as follows: magenta, Lhcq subfamily; red, Lhcz subfamily; orange, Lhcr subfamily; brown, CgLhcr9 homologs; green, Lhcf subfamily (CgLhcf9 homolog clade is in gray); blue, Lhcx subfamily. The following diatom species were analyzed: **A**, *Fistulifera solaris*; **B**, *Fragilariopsis cylindrus*; **C**, *Pseudo-nitzschia multistriata*; **D**, *Thalassiosira oceanica*.

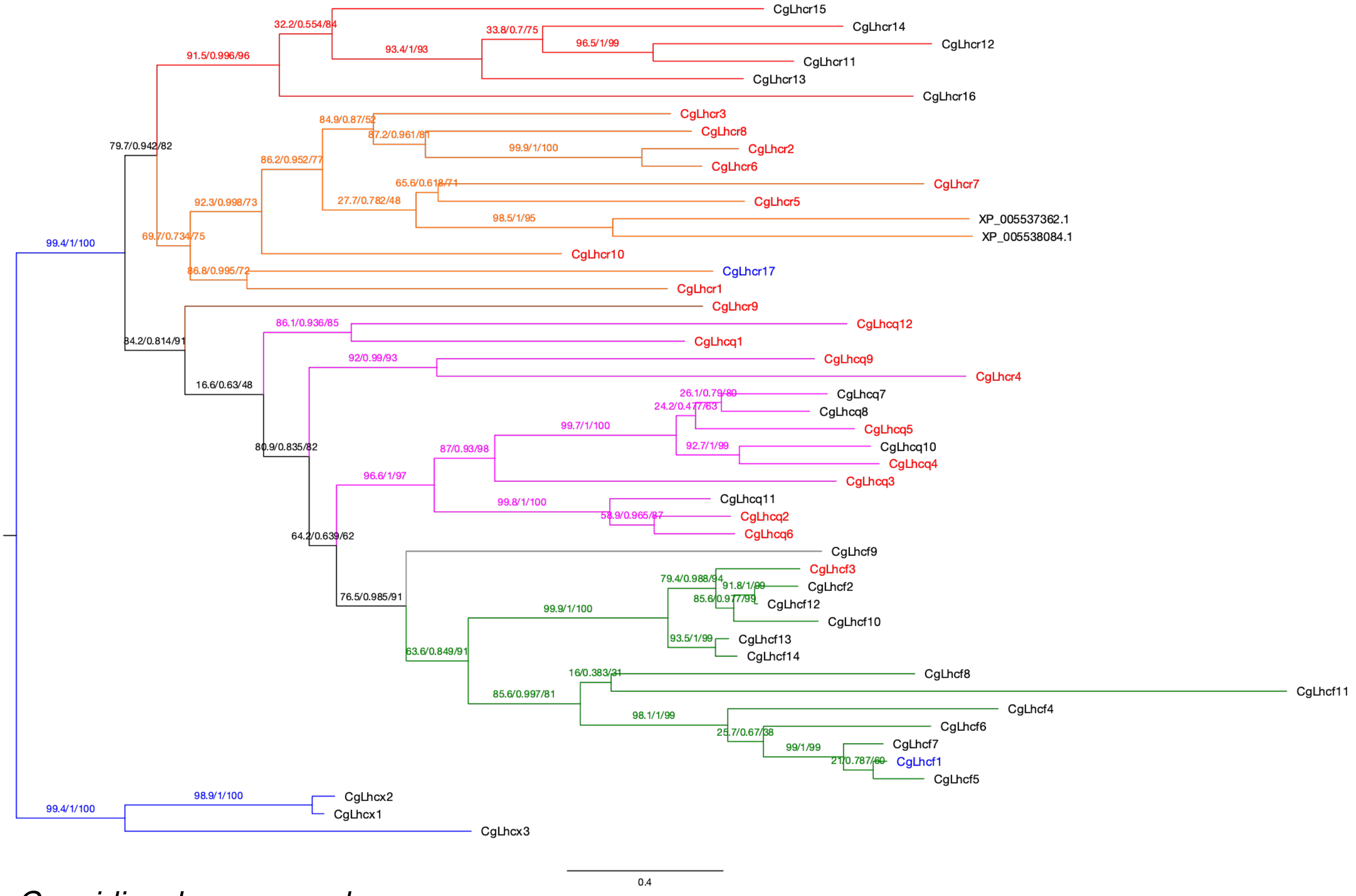


*Cyanidioschyzon merolae*

Supplemental Figure S3A

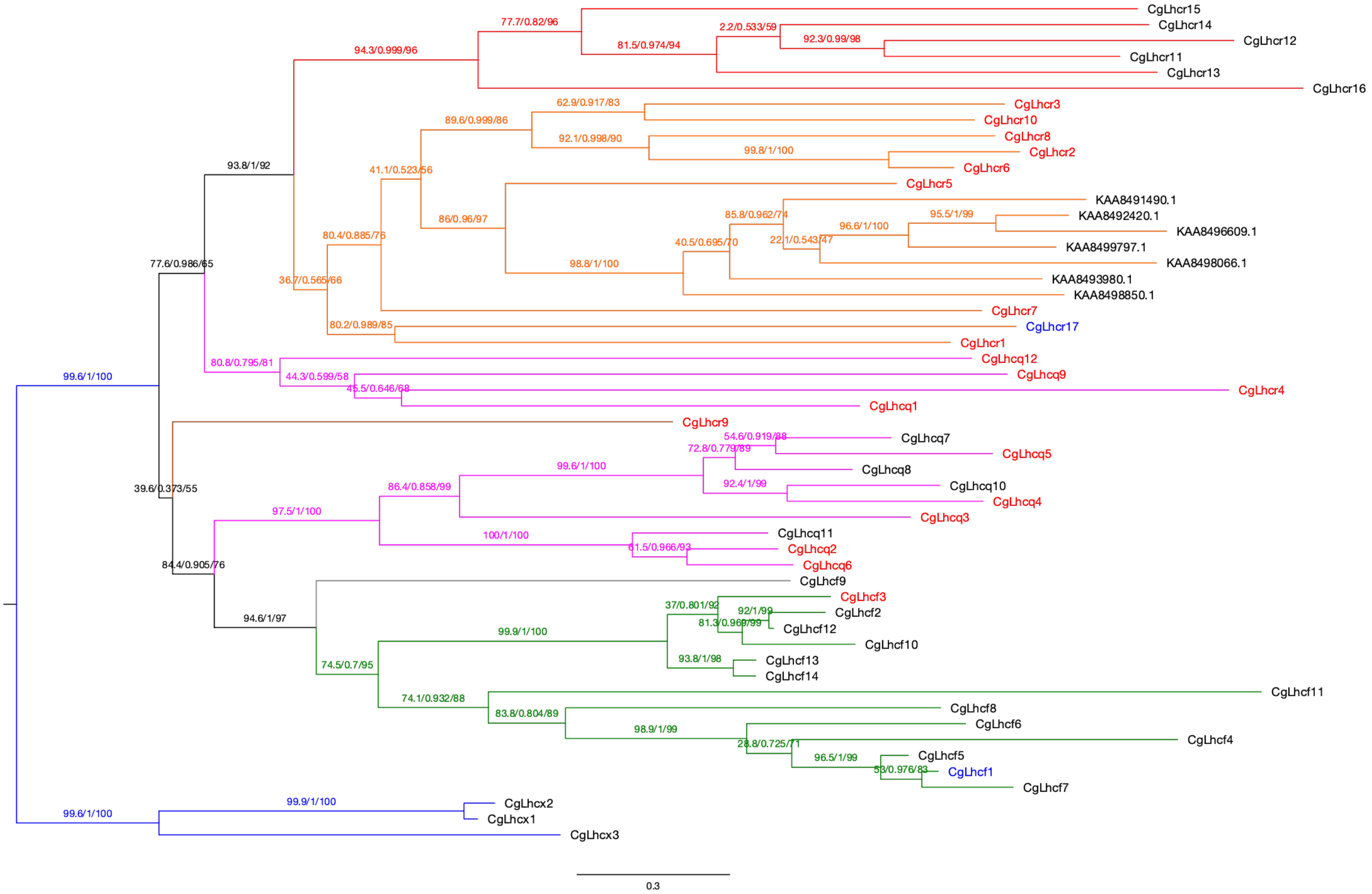


*Porphyridium purpureum*

Supplemental Figure S3B

**Supplemental Figure S3. Maximum likelihood tree of FCPs/LHCs from *Chaetoceros gracilis* and red algae.** The trees were inferred using IQ-TREE 2 (Minh *et al.*, 2020). Each gene is labeled with NCBI RefSeq or GenBank accession ID. The numbers of supporting values are SH-aLRT support (%)/aBayes support/ultrafast bootstrap support (%). Colors of clades are as follows: magenta, Lhcq subfamily; red, Lhcz subfamily; orange, Lhcr subfamily; brown, CgLhcr9 homologs; green, Lhcf subfamily (CgLhcf9 homolog clade is in gray); blue, Lhcx subfamily. The following red-algal species were analyzed: **A,** *Cyanidioschyzon merolae*; **B,** *Porphyridium purpureum*.

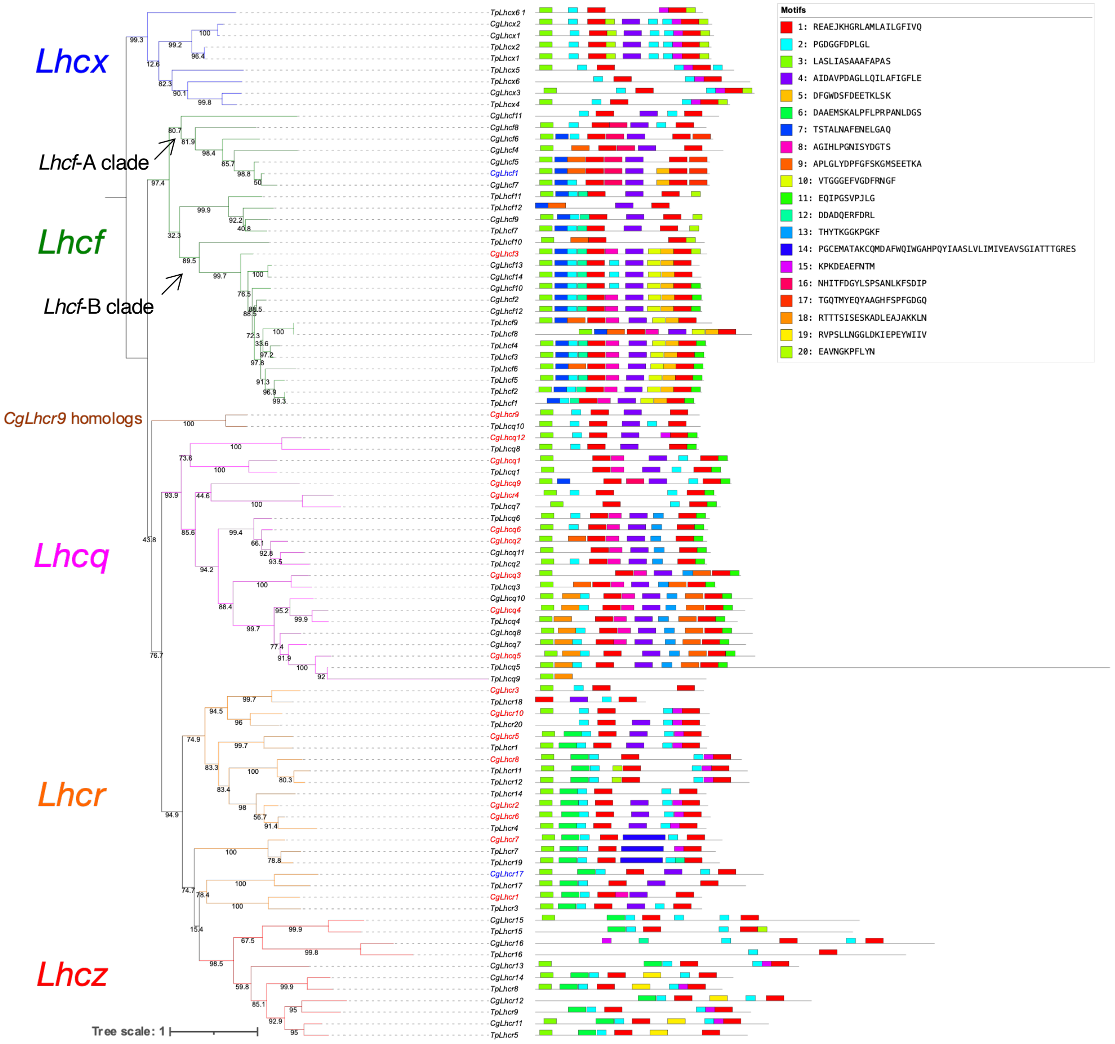


**Supplemental Figure S4. Maximum-likelihood tree of FCPs from *Chaetoceros gracilis* and *Thalassiosira pseudonana* showing the localization of the motifs generated by MEME.** Multiple Expectation maximizations for Motif Elicitation (MEME, version 5.3.0) (Bailey *et al.*, 2009) was performed using translated sequences of FCP genes from *Chaetoceros gracilis* and *Thalassiosira pseudonana.* The tree was rerooted with the Lhcx subfamily. The numbers of supporting values are ultrafast bootstrap support (%). Colors of clades are as follows: magenta, Lhcq subfamily; red, Lhcz subfamily; orange, Lhcr subfamily; brown, CgLhcr9 homologs; green, Lhcf subfamily; blue, Lhcx subfamily. Colors of gene names are as follows: red, PSI-assigned FCP; blue PSII-assigned FCP. In this figure, Lhcf subfamily has 3 clades: CgLhcf9 homologs, Lhcf-A and Lhcf-B.

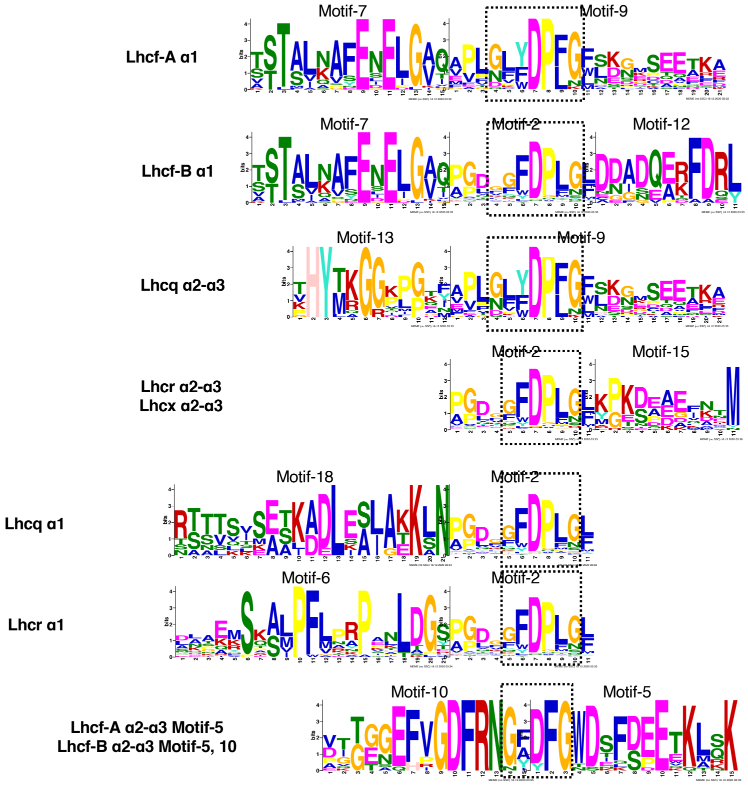


**Supplemental Figure S5. MEME motif logos contained the conserved carotenoid-binding motif "GFDPLG" with adjacent MEME motif logos.** The motifs were generated from *Chaetoceros gracilis* and *Thalassiosira pseudonana* FCP amino acid sequences using MEME (Bailey *et al.*, 2009). The region of the Car-binding motif in each FCP/LHC subfamily are indicated as dashed squares. " Lhcf-A" and "Lhcf-B" corresponds to the clades indicated in Supplemental Fig. S4.

B

```
--------------------------------------------------------------------------------
         Motif EQIPGSVPJLG MEME-11 sites sorted by position p-value
--------------------------------------------------------------------------------
Sequence name            Start   P-value                  Site
-------------            ----- ---------             -----------
TpLhcq2                    191  1.31e-11 MIGMMSFIAA HNIPGSVPVLG SNF
CgLhcr4                    202  8.48e-11 MIAMASLFAM EAIPGSVPIMN VFN
CgLhcq7                    220  2.91e-10 QLGIIGFLCA QTIPGSVPALV GVVKPYSGEV
CgLhcq5                    227  3.30e-10 MLGIFGFLCA DTIPGSVPLLN DIAIPYSGQV
CgLhcq8                    223  3.76e-10 MLGLFGFLCA QTIPGSVPLLS GVLPAYDGEP
CgLhcf10                   186  4.80e-10 MFGILGLMVH EQLGGSLPIVG EM
CgLhcq10                   223  1.07e-09 MLGIFGFISE QKIEGSVPLLK GVVPAYDGEF
TpLhcq1                    210  1.67e-09 MITMAGWLSE ESIPGSFPVWH P
CgLhcq2                    189  1.67e-09 MIAIMSFISA ANIPGSVPALT NNPAF
TpLhcf4                    192  1.86e-09 MMGILGLMMH EQLGGSIPIVG EL
CgLhcq3                    232  1.86e-09 MIGVAGLYCA ATIPGSVPLQP PC
CgLhcq11                   193  1.86e-09 MIGIMSFIAA NNIPGSVPALV GNPAF
TpLhcq6                    192  2.57e-09 MISIMSFVAA ANIPGSVPALA GNPHF
CgLhcq6                    190  2.57e-09 MIAIMSFVAA ANIPGSVPALA GNPMF
CgLhcf12                   186  2.85e-09 MMGILGLMVH EQLGGTLPIVG EM
CgLhcq9                    221  2.85e-09 MFAISAWVAA EAIPGSVPIPL PW
CgLhcf2                    187  2.85e-09 MMGILGLMVH EQLGGTLPIVG EM
TpLhcq7                    205  3.49e-09 MIAMASLFAM KAIPGSVPIMD MLGAN
TpLhcf6                    189  4.67e-09 MMGILGLMMH EQVGGTFPIVG EM
TpLhcq8                    181  5.66e-09 MIGIAGFAAN HFIPGSCPVPD FIA
TpLhcf1                    178  8.23e-09 MMGILGLMVH EQLGGELPIVG QM
TpLhcf2                    186  8.23e-09 MMGILGLMVH EQLGGELPIVG QM
TpLhcf5                    188  8.23e-09 MMGILGLMVH EQLGGELPIVG QM
CgLhcq4                    224  8.23e-09 MIGIMGFLAE QKVEGSVPLLK GVVPHYDGEV
TpLhcq5                    218  1.55e-08 QLGIFGFLCA DKITGSVPALN SIAIHYDGNP
TpLhcq4                    215  1.69e-08 MIGIMGFISE QRVEGSVPLLK GLVAHYDGET
CgLhcq1                    218  2.37e-08 MMTIAAWVSD EMIPGALPVWH P
TpLhcq3                    203  2.79e-08 MIGVAGMYAA ATIDGSVPLQP PC
TpLhcf3                    190  3.03e-08 MMGILGLMMH EQIGGVFPITG EL
CgLhcq12                   182  5.72e-08 MLGIAGFVSA HFIPGAVPIGP GF
```

A

Car binding motif

```
CgLhcq7  K-KLNPIVG---------YFDPLG
CgLhcq8  K-KLNPIVG---------FYDPLN
CgLhcq5  K-KANPTLG---------YYDPLS
TpLhcq5  T-KLNPTIR---------FYDPLN
CgLhcq10 K-KLNPVVP---------RFDPLG
CgLhcq4  E-KSNPVLK---------YYDPLQ
TpLhcq4  E-KSNPVLK---------FYDPLE
CgLhcq11 G-ATAP-LK---------NFDPLR
CgLhcq2  G-STAP-VG---------PFDPLN
CgLhcq6  G-ATAP-LG---------KFDPLN
TpLhcq6  G-ATAP-LK---------GFDPLN
TpLhcq2  G-ASAP-LK---------NFDPLN
TpLhcq9  T-KLNPTIR---------FYDPLN

CgLhcq12 G-TSVE-TG-------GKIFDPLN
CgLhcq9  --GATP-LGGEPVFIGENYWDKLT
CgLhcr4  G----------------KPWDPLS
TpLhcq7  G----------------EVFDPMG

CgLhcq3  S-PDTTEIG--------RVWDPLG
TpLhcq3  E-PDTVEFG--------QKWDPLG
CgLhcq1  G-ISIE-TG--------NKCPPLG
TpLhcq1  G-VTNE-TG--------NKIPPLG
```

C

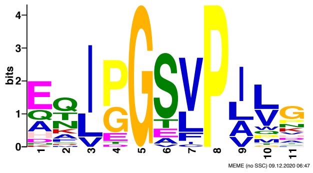


**Supplemental Figure S6. Specific motifs of Lhcq subfamily proteins: varieties of carotenoid-binding motifs and the novel C-terminal chlorophyll binding motif "PGSVP". A,** The multiple alignment of Car-binding motif of Lhcq subfamily proteins. **B,** MEME report of the novel conserved chlorophyll binding motif "PGSVP". **C,** The logo of the MEME-11 motif containing "PGSVP".

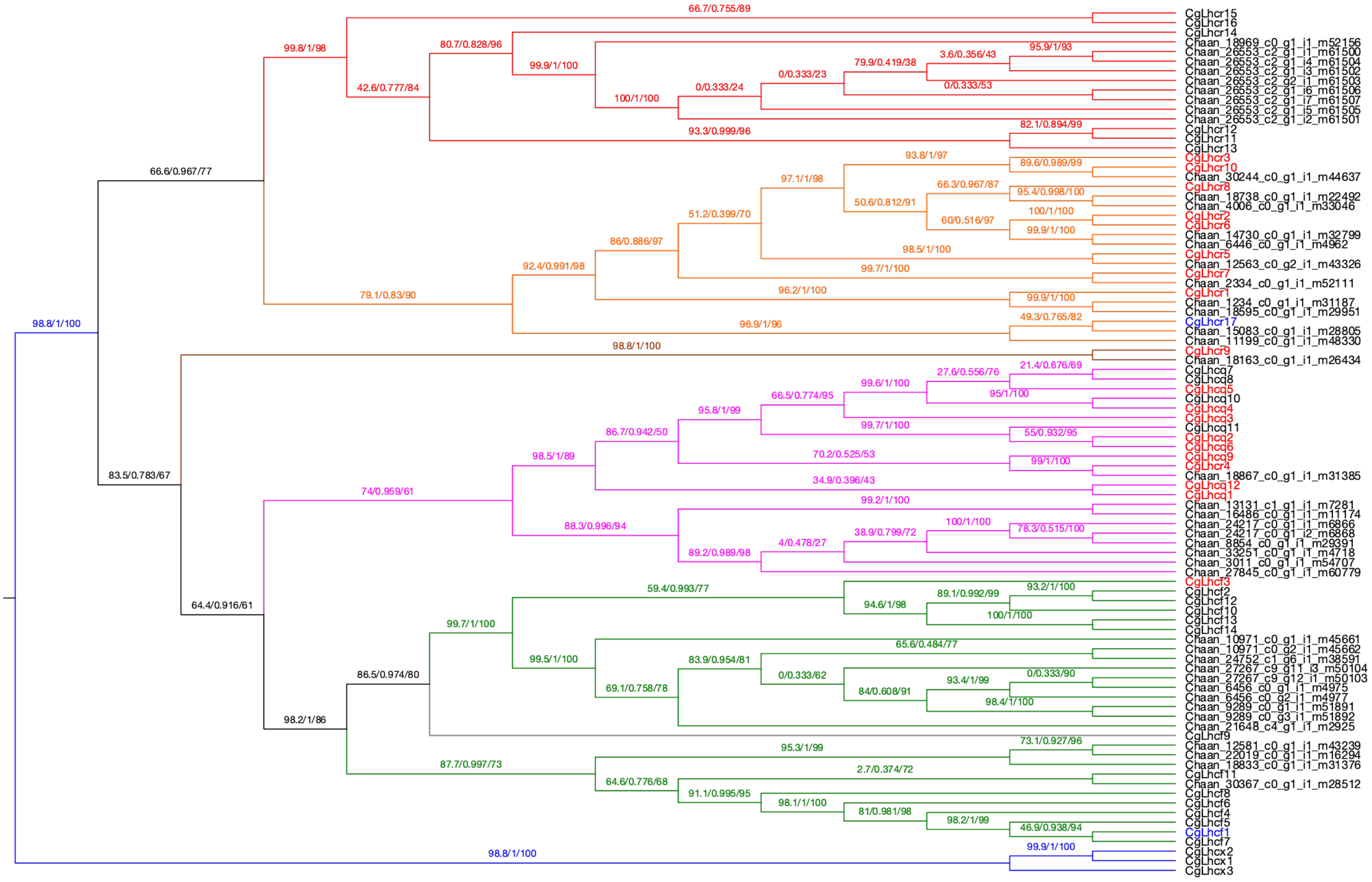


*Chattonella antiqua*

Supplemental Figure S7A

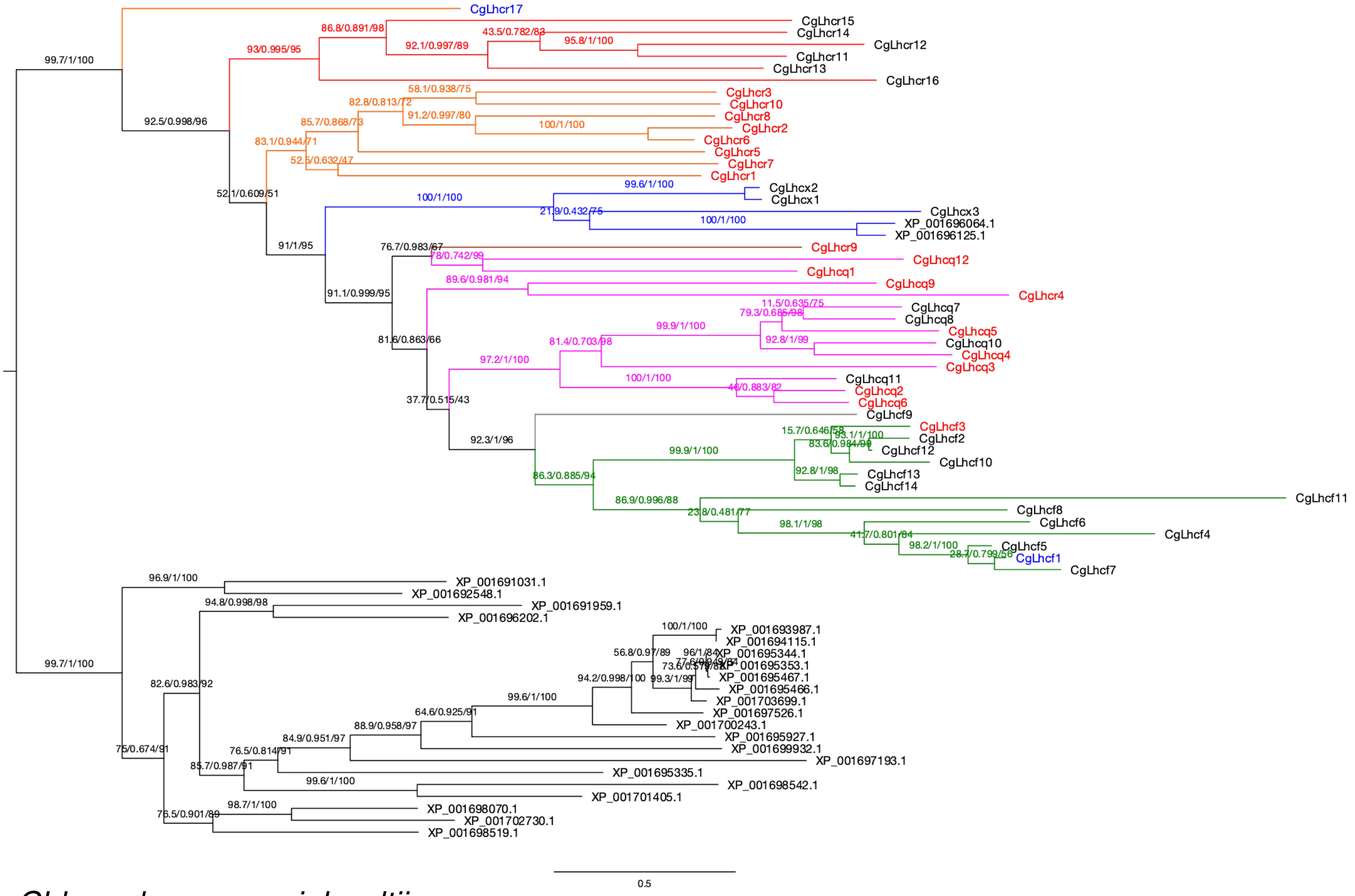


*Chlamydomonas reinhardtii*

Supplemental Figure S7B

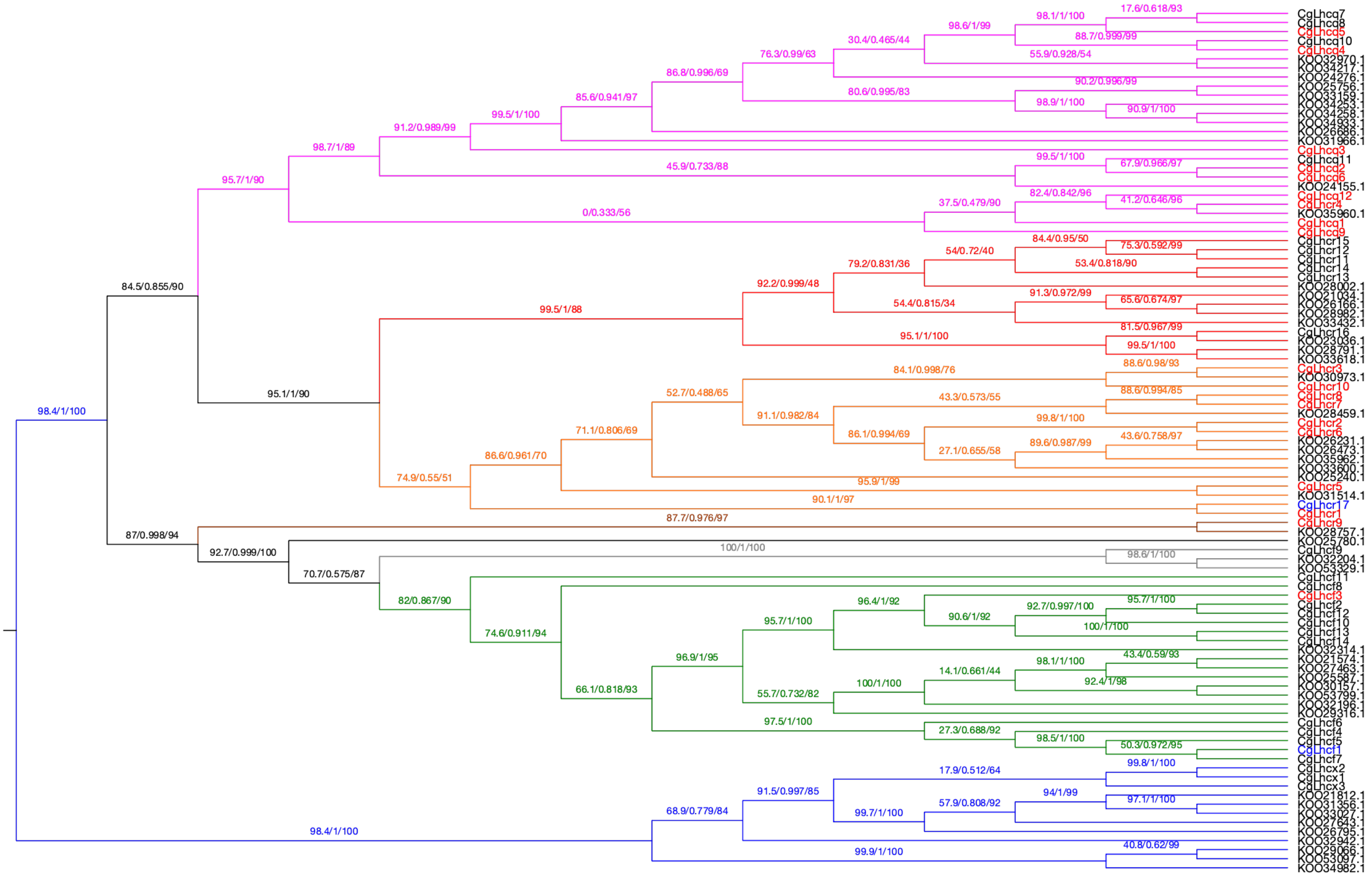


*Chrysochromulina tobinii*

Supplemental Figure S7C

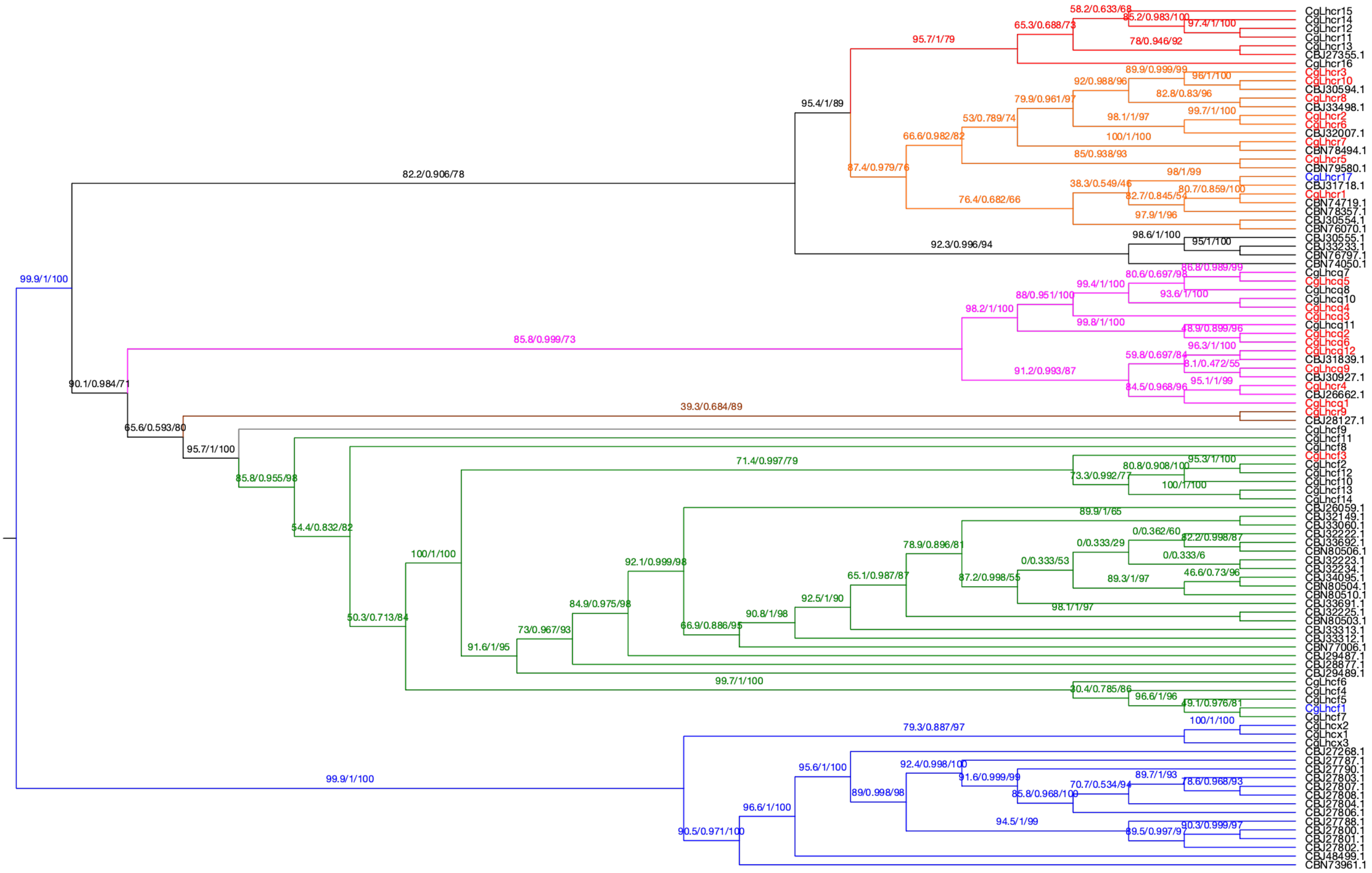

*Ectocarpus siliculosus*

Supplemental Figure S7D

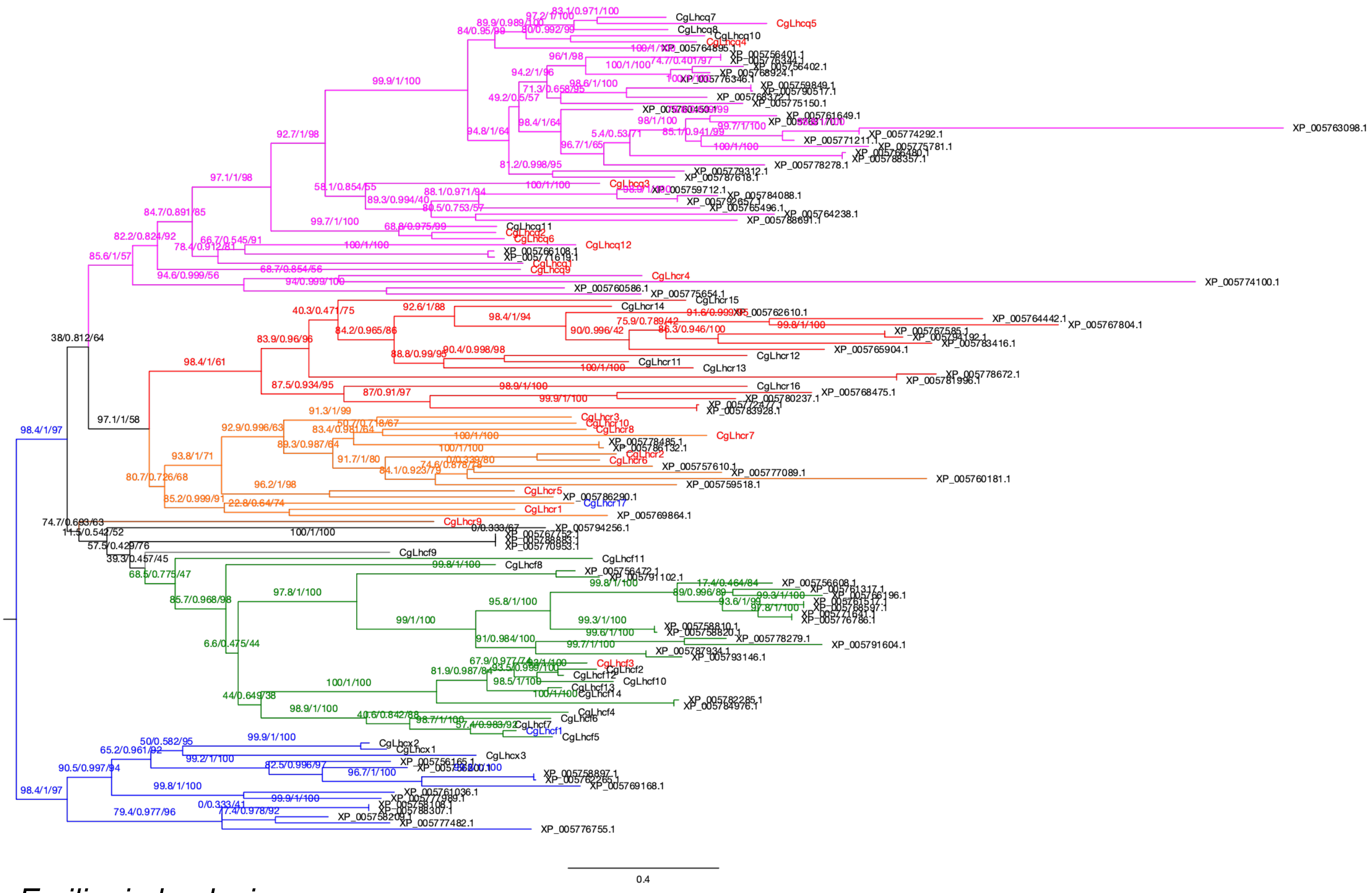


*Emiliania huxleyi*

Supplemental Figure S7E

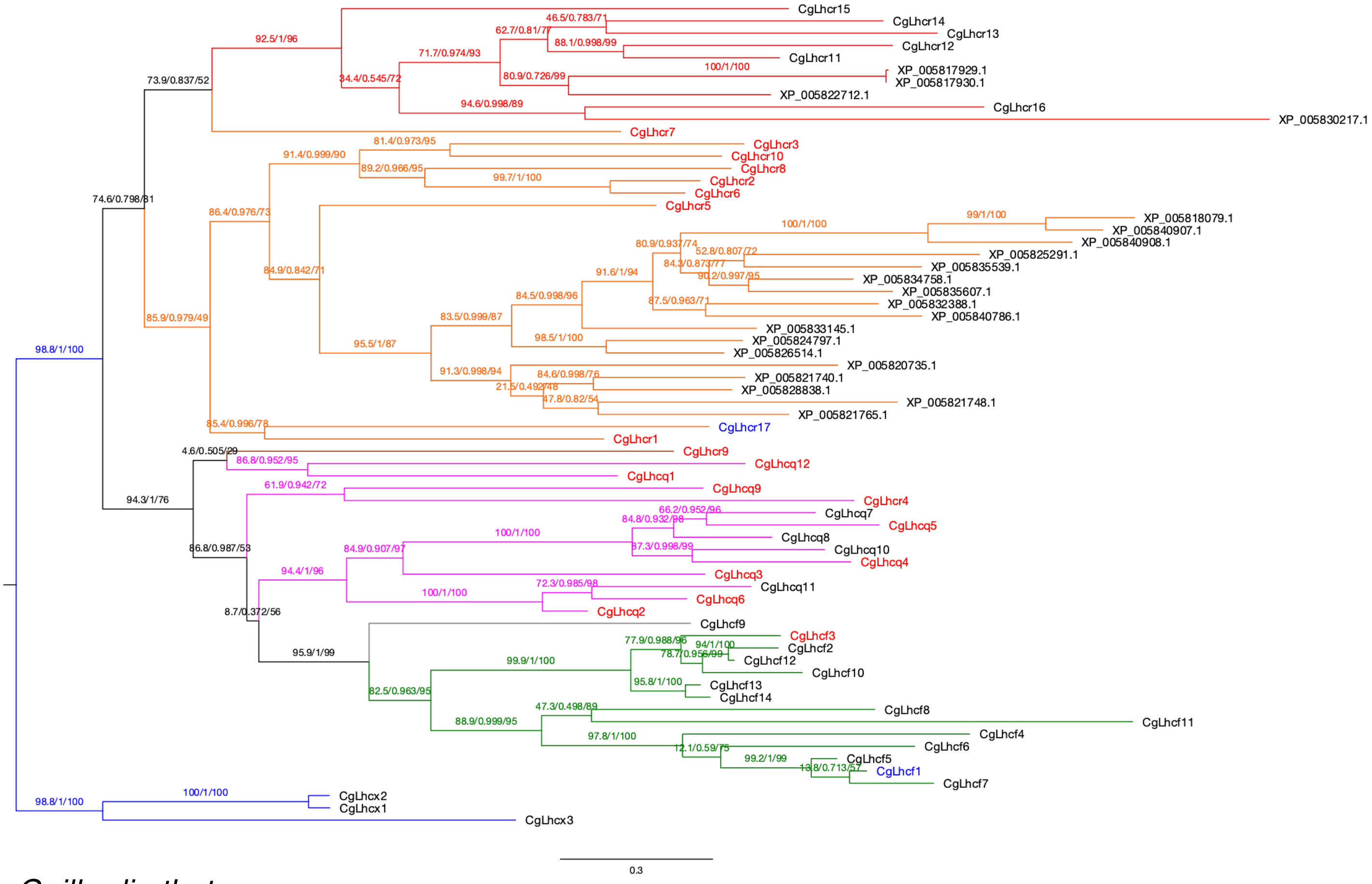


*Guillardia theta*

Supplemental Figure S7F

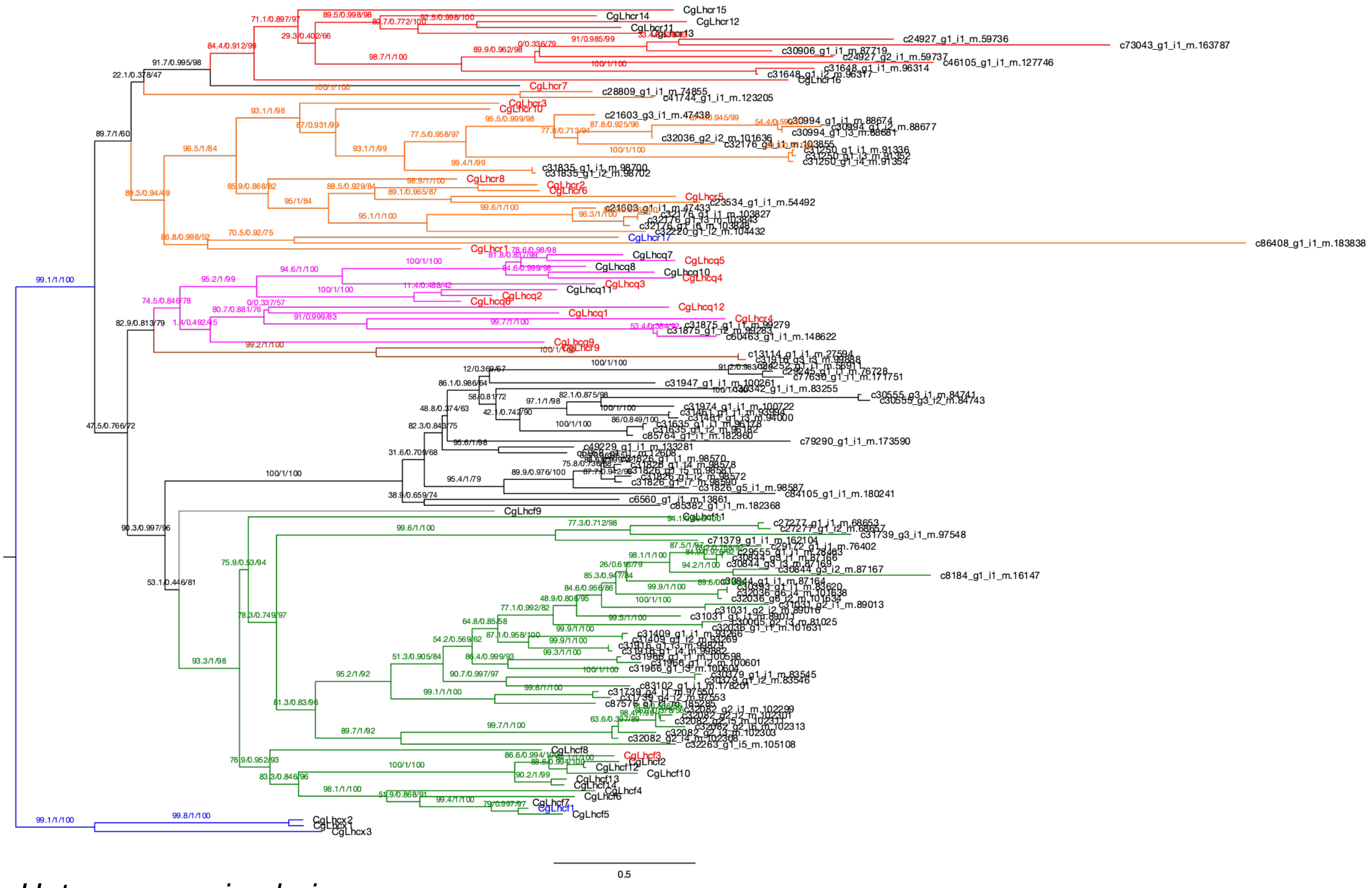

*Heterocapsa circularisquama*

Supplemental Figure S7G

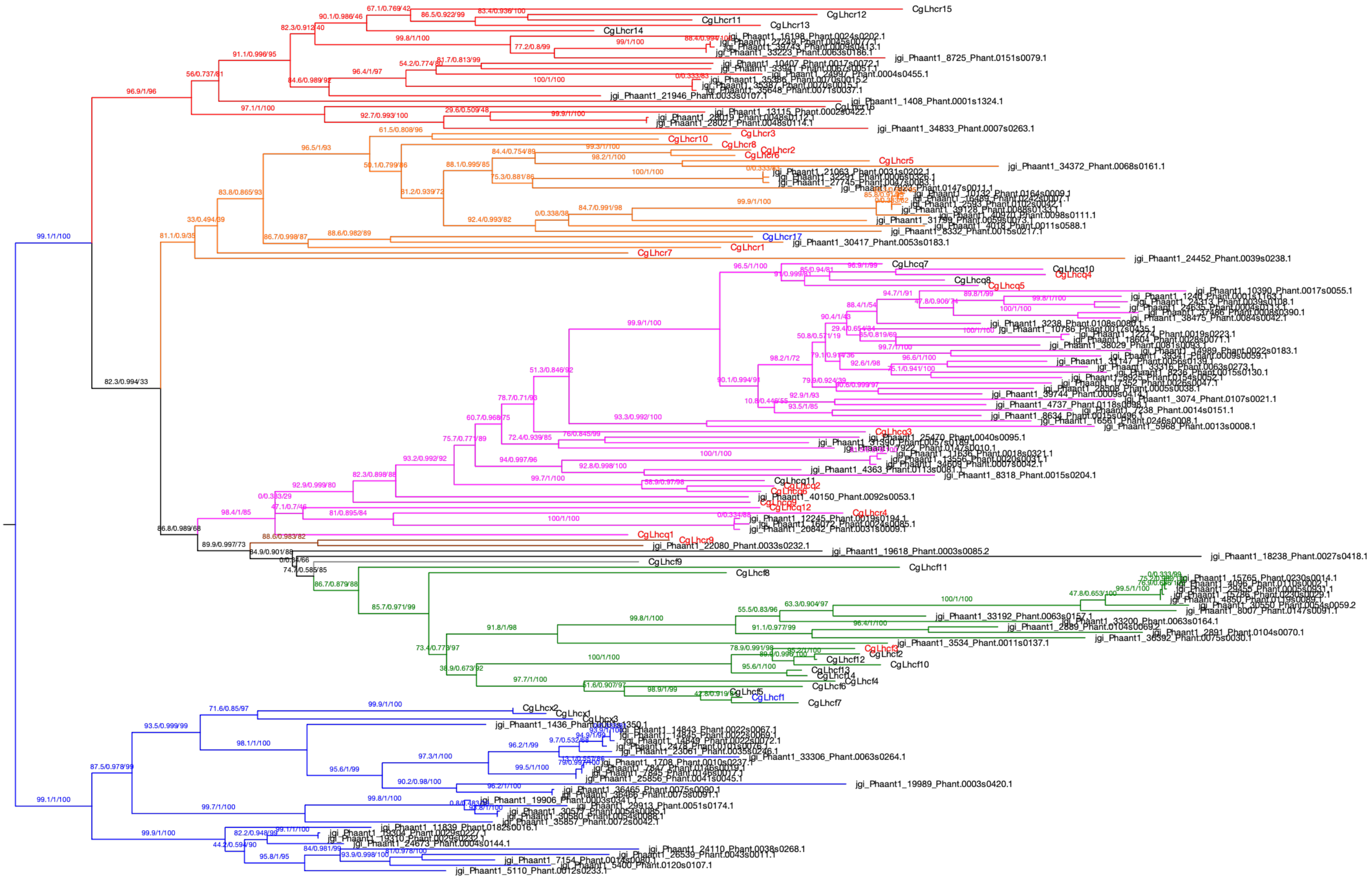

*Phaeocystis antarctica*

Supplemental Figure S7H

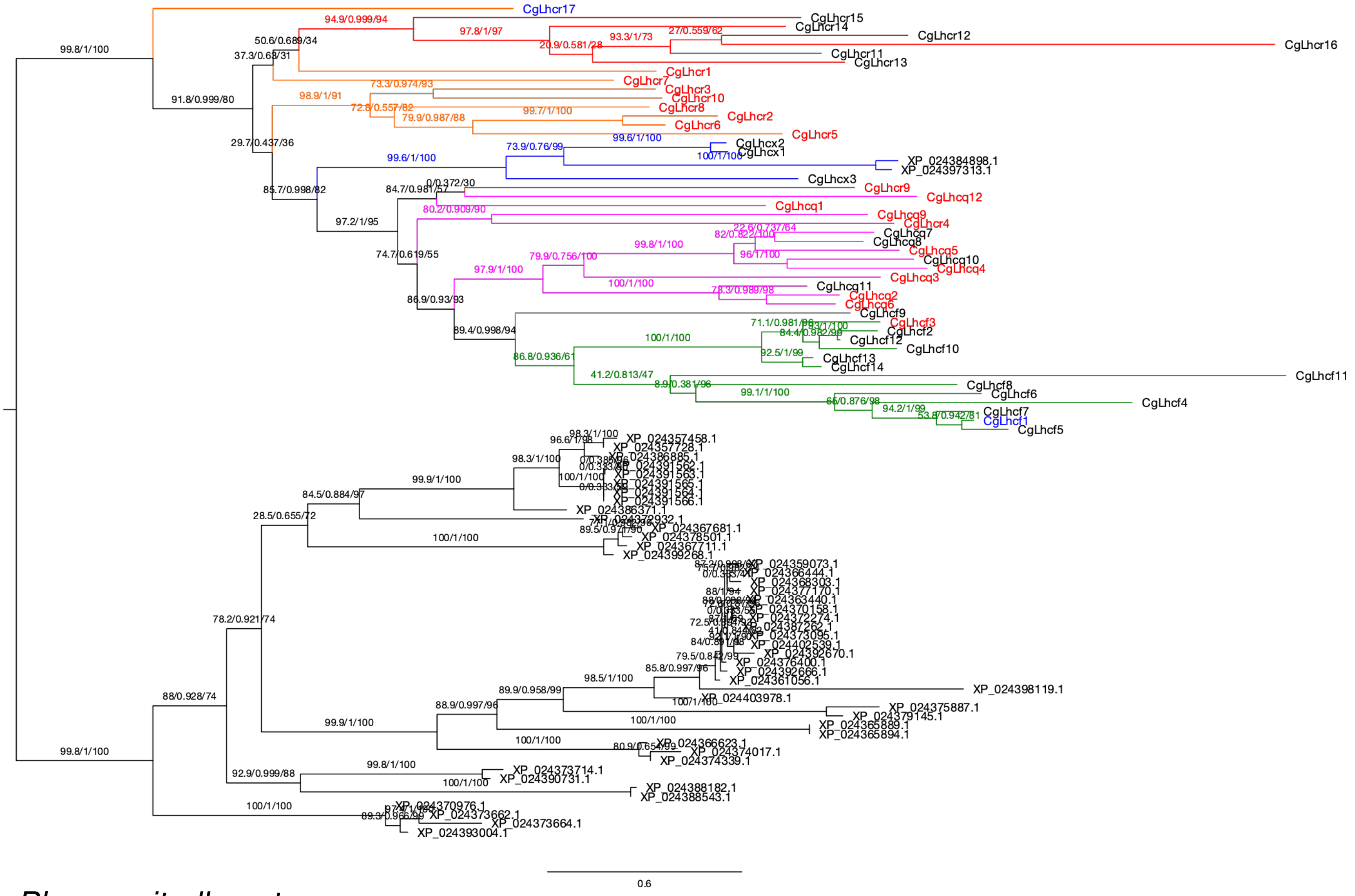


*Physcomitrella patens*

Supplemental Figure S7I

**Supplemental Figure S7. Maximum-likelihood phylogenetic tree of FCPs/LHCs from *Chaetoceros gracilis* and red- and green-lineage species.** The numbers of supporting values are SH-aLRT support (%)/aBayes support/ultrafast bootstrap support (%). Colors of clades are as follows: magenta, Lhcq subfamily; red, Lhcz subfamily; orange, Lhcr subfamily; brown, CgLhcr9 homologs; green, Lhcf subfamily (CgLhcf9 homolog clade is in gray); blue, Lhcx subfamily. The species used in the tree are mentioned below.

**A,** *Chattonella antiqua*

**B,** *Chlamydomonas reinhardtii*

**C,** *Chrysochromulina tobinii*

**D,** *Ectocarpus siliculosus*

**E,** *Emiliania huxleyi*

**F,** *Guillardia theta*

**G,** *Heterocapsa circularisquama*

**H,** *Phaeocystis antarctica*

**I,** *Physcomitrella patens*

**Supplemental Table S1. List of assemblies used in FCP/LHC detection with BUSCO scores and lineages.** Benchmarking Universal Single-Copy Orthologs (BUSCO v4.0.6, Seppey et al. 2019) was performed in protein mode. BUSCO lineage datasets used in our analyses were selected based on their taxonomy listed in the table.

| Species | Assembly-type | BUSCO v4.0.6 | BUSCO lineage |
|---|---|---|---|
| *Cyanidioschyzon merolae* | Genome | 72.6% | eukaryota_odb10 |
| *Porphyridium purpureum* | Genome | 72.6% | eukaryota_odb10 |
| *Ectocarpus siliculosus* | Genome | 97% | stramenopiles_odb10 |
| *Chattonella antiqua* | Transcriptome | 99% | stramenopiles_odb10 |
| *Heterocapsa circularisquama* | Transcriptome | 84.2% | alveolata_odb10 |
| *Pseudo-nitzschia multistriata* | Genome | 86% | stramenopiles_odb10 |
| *Fragilariopsis cylindrus* | Genome | 95% | stramenopiles_odb10 |
| *Fistulifera solaris* | Genome | 97% | stramenopiles_odb10 |
| *Phaeodactylum tricornutum* | Genome | 97% | stramenopiles_odb10 |
| *Chaetoceros gracilis* | Genome | 96% | stramenopiles_odb10 |
| *Thalassiosira pseudonana* | Genome | 97% | stramenopiles_odb10 |
| *Thalassiosira oceanica* | Genome | 90% | stramenopiles_odb10 |
| *Emiliania huxleyi* | Genome | 56.1% | eukaryota_odb10 |
| *Chrysochromulina tobinii* | Genome | 53.0% | eukaryota_odb10 |
| *Phaeocystis antarctica* | Genome | 70.2% | eukaryota_odb10 |
| *Chlamydomonas reinhardtii* | Genome | 93.0% | chlorophyta_odb10 |
| *Physcomitrella patens* | Genome | 88.8% | embryophyta_odb10 |
| *Guillardia theta* | Genome | 79.6% | eukaryota_odb10 |

**Supplemental Table S2. List of *Chaetoceros gracilis* FCPs with gene IDs or accession IDs.**

| Gene name | ID |
|---|---|
| CgLhcf1 | g14370.t1 |
| CgLhcf5 | g2935.t1 |
| CgLhcf7 | g6797.t1 |
| CgLhcf6 | g5499.t1 |
| CgLhcf2 | g4294.t1 |
| CgLhcf13 | LC647435 |
| CgLhcf14 | LC647436 |
| CgLhcf12 | g14457.t1 |
| CgLhcf3 | g886.t1 |
| CgLhcf10 | g8542.t1 |
| CgLhcf4 | g2721.t1 |
| CgLhcf8 | g7443.t1 |
| CgLhcx1 | g4948.t1 |
| CgLhcx2 | g1219.t1 |
| CgLhcf9 | g7689.t1 |
| CgLhcx3 | g8495.t1 |
| CgLhcr9 | g2996.t1 |
| CgLhcq7 | g156.t1 |
| CgLhcq8 | g2936.t1 |
| CgLhcq5 | g4794.t1 |
| CgLhcq4 | g7567.t1 |
| CgLhcq10 | g987.t1 |
| CgLhcq11 | g857.t1 |
| CgLhcq2 | g858.t1 |
| CgLhcq6 | g2709.t1 |
| CgLhcq3 | g7730.t1 |
| CgLhcf11 | g9284.t1 |
| CgLhcq1 | g11462.t1 |
| CgLhcq12 | g4203.t1 |
| CgLhcr3 | g1479.t1 |
| CgLhcr10 | g1480.t1 |
| CgLhcr8 | g2857.t1 |
| CgLhcr2 | g10324.t1 |
| CgLhcr6 | g11475.t1 |
| CgLhcr5 | g11413.t1 |
| CgLhcr1 | g11656.t1 |
| CgLhcr14 | g3776.t1 |
| CgLhcr12 | g5385.t1 |
| CgLhcr11 | g11554.t1 |
| CgLhcr13 | g11553.t1 |
| CgLhcr7 | g11027.t1 |
| CgLhcr15 | g1057.t1 |
| CgLhcq9 | g9170.t1 |
| CgLhcr17 | g3978.t1 |
| CgLhcr4 | g10543.t1 |
| CgLhcr16 | g10173.t1 |

**Supplemental Table S3. List of all FCPs from *Thalassiosira pseudonana* with revised gene names.** The columns: "Former Gene names", "Cross-refference (RefSeq)" and "Protein names" were downloaded from Uniprot (https://www.uniprot.org/). New gene names written in bold are renamed FCPs from former names.

| New Gene Names | Former Gene names | Cross-reference (RefSeq) | Protein names |
|---|---|---|---|
| TpLhcr9 | Lhcr9 THAPSDRAFT_23889 | XP_002292153.1; | Fucoxanthin chl a/c light-harvesting protein |
| TpLhcr1 | Lhcr1 THAPSDRAFT_24080 | XP_002292353.1; | Fucoxanthin chl a/c light-harvesting protein, lhcr type |
| TpLhcr7 | Lhcr7 THAPSDRAFT_3816 | XP_002288517.1; | Fucoxanthin chl a/c light-harvesting protein |
| TpLhcr5 | Lhcr5 THAPS_25433 | XP_002295649.1; | Fucoxanthin chlorophyll a/c light-harvesting protein |
| TpLhcr4 | Lhcr4 THAPSDRAFT_4882 | XP_002289997.1; | Fucoxanthin chl a/c light-harvesting protein |
| TpLhcr11 | Lhcr11 THAPSDRAFT_2342 | XP_002287251.1; | Fucoxanthin chlorophyll a/c light-harvesting protein |
| TpLhcr3 | Lhcr3 THAPSDRAFT_26246 | XP_002290650.1; | Fucoxanthin chl a/c light-harvesting protein |
| **TpLhcq7** | Lhcr10 THAPSDRAFT_22747 | XP_002290280.1; | Fucoxanthin chlorophyll a/c light-harvesting protein |
| TpLhcf7 | Lhcf7 THAPSDRAFT_21667 | XP_002288232.1; | Fucoxanthin chlorophyll a/c light-harvesting protein, major type |
| TpLhcr14 | Lhcr14 THAPSDRAFT_4883 | XP_002289542.1; | Fucoxanthin chlorophyll a/c light-harvesting protein, lhcr type |
| TpLhcf10 | Lhcf10 THAPS_25402 | XP_002295619.1; | Fucoxanthin chl a/c light-harvesting protein, major type |
| TpLhcr8 | Lhcr8 THAPSDRAFT_11272 | XP_002297122.1; | Fucoxanthin chl a/c light-harvesting protein |
| **TpLhcr20** | lhcA THAPSDRAFT_bd218 | XP_002297311.1; | Photosystem I light harvesting protein (Fragment) |
| NA | 3HfcpB | | Fucoxanthin-chlorophyll a/c light-harvesting protein (Fragment) |
| NA | 3HfcpA | | Fucoxanthin-chlorophyll a/c light-harvesting protein (Fragment) |
| **TpLhcr16** | THAPSDRAFT_4871 | XP_002289537.1; | Uncharacterized protein |
| **TpLhcq5** | THAPSDRAFT_10219 | XP_002294116.1; | Uncharacterized protein |
| **TpLhcr17** | Lhca2 THAPSDRAFT_2601 | XP_002287377.1; | Fucoxanthin chl a/c protein, lhca clade |
| **TpLhcr19** | Lhca6 THAPSDRAFT_3815 | XP_002289005.1; | Member of fucoxanthin chlorophyll a/c family, lhca clade |
| TpLhcx1 | lhcx1 THAPSDRAFT_264921 | XP_002295258.1; | Fucoxanthin chlorophyll a/c protein, LI818 clade |
| TpLhcx2 | Lhcx2 THAPSDRAFT_38879 | XP_002295184.1; | Fucoxanthin chlorophyll a/c protein, LI818 clade |
| TpLhcx6 | Lhcx6 THAPSDRAFT_12097 | XP_002295183.1; | Fucoxanthin chlorophyll a/c protein, LI818 clade |
| TpLhcx6_1 | Lhcx6_1 THAPS_30385 | XP_002295578.1; | Fucoxanthin chlorophyll a/c protein-LI818 clade |
| TpLhcr12 | Lhcr12 THAPSDRAFT_2341 | XP_002287768.1; | Fucoxanthin chlorophyll a/c protein |
| **TpLhcr15** | FCP_1 THAPSDRAFT_270220 | XP_002295168.1; | Uncharacterized protein |
| **TpLhcq1** | FCP2 THAPSDRAFT_4843 | XP_002289522.1; | Uncharacterized protein |
| **TpLhcq2** | FCP_2 THAPSDRAFT_7916 | XP_002292206.1; | Uncharacterized protein |
| TpLhcf6 | Lhcf6 THAPSDRAFT_33018 | XP_002288729.1; | Fucoxanthin chlorophyll a/c protein 6 |
| TpLhcx5 | lhcx5 THAPSDRAFT_31128 | XP_002287075.1; | Fucoxanthin chlorophyll a/c protein, LI818 clade |
| TpLhcf9 | Lhcf9 THAPSDRAFT_268127 | XP_002286436.1; | Fucoxanthin chlorophyll a/c protein 8 |
| **TpLhcq10** | FCP3 THAPSDRAFT_11501 | XP_002294580.1; | Pt17531-like protein |
| TpLhcf4 | Lhcf4 THAPSDRAFT_38667 | XP_002294845.1; | Fucoxanthin chlorophyll a/c protein 4 |
| TpLhcf5 | Lhcf5 THAPSDRAFT_42962 | XP_002294608.1; | Fucoxanthin chlorophyll a/c protein 5 |
| TpLhcf11 | Lhcf11 THAPSDRAFT_270241 | XP_002288023.1; | Fucoxanthin-chlorophyll a-c binding protein, plastid |
| TpLhcx4 | Lhcx4 THAPSDRAFT_270228 | XP_002290755.1; | Fucoxanthin chlorophyll a/c protein, LI818 clade |
| **TpLhcq3** | THAPSDRAFT_6139 | XP_002291432.1; | Uncharacterized protein |
| **TpLhcq6** | THAPSDRAFT_270221 | XP_002288721.1; | Uncharacterized protein |
| **TpLhcf3** | THAPSDRAFT_270233 | XP_002295015.1; | Uncharacterized protein |
| **TpLhcq4** | THAPSDRAFT_270092 | XP_002294886.1; | Uncharacterized protein |
| **TpLhcq8** | THAPSDRAFT_23808 | XP_002292077.1; | Uncharacterized protein |
| TpLhcf8 | Lhcf8 THAPSDRAFT_5174 | XP_002290137.1; | Fucoxanthin chlorophyll a/c protein 8 |
| TpLhcf2 | Lhcf2 THAPSDRAFT_260392 | XP_002294743.1; | Fucoxanthin chlorophyll a/c protein 2 (Fragment) |
| TpLhcf1 | Lhcf1 THAPSDRAFT_38583 | XP_002295087.1; | Fucoxanthin chlorophyll a/c protein 1 (Fragment) |
| **TpLhcr18** | THAPSDRAFT_bd1160 | XP_002297322.1; | Uncharacterized protein (Fragment) |
| **TpLhcf12** | THAPSDRAFT_264714 | XP_002294610.1; | Fucoxanthin chl a/c protein-deviant sequence (Fragment) |
| **TpLhcq9** | THAPSDRAFT_270329 | XP_002294117.1; | Uncharacterized protein |

**Supplemental Table S4. List of all FCPs from *Phaeodactylum tricornutum* with revised gene names.** The columns: "Former Gene names", "Cross-refference (RefSeq)" and "Protein names" were downloaded from Uniprot (https://www.uniprot.org/). New gene names written in bold are renamed FCPs from former names.

| New Gene Names | Former Gene Names | Cross-reference (RefSeq) | Protein names |
|---|---|---|---|
| **PtLhcr16** | PHATRDRAFT_6062 | XP_002182909.1; | Predicted protein (Fragment) |
| **PtLhcf18** | PHATRDRAFT_56448 | XP_002177056.1; | Fucoxanhin chlorophyll binding protein related |
| **PtLhcq1** | PHATRDRAFT_48798 | XP_002183454.1; | Fucoxanthin chlorophyll a/c protein, deviant |
| **PtLhcq2** | PHATRDRAFT_47485 | XP_002181795.1; | Fucoxanthin chlorophyll a/c protein, deviant (Fragment) |
| **PtLhcq3** | PHATRDRAFT_24119 | XP_002185437.1; | Fucoxanthin chlorophyll a/c protein, deviant |
| **PtLhcq5** | PHATRDRAFT_17531 | XP_002176735.1; | Fucoxanthin chlorophyll a/c protein |
| **PtLhcr15** | PHATRDRAFT_15820 | XP_002183911.1; | Predicted protein (Fragment) |
| PtLhcx4 | Lhcx4 PHATRDRAFT_38720 | XP_002182760.1; | Protein fucoxanthin chlorophyll a/c protein |
| PtLhcx3 | Lhcx3 PHATRDRAFT_44733 | XP_002178699.1; | Protein fucoxanthin chlorophyll a/c protein |
| PtLhcx2 | Lhcx2 PHATRDRAFT_54065 | XP_002176987.1; | Protein fucoxanthin chlorophyll a/c protein |
| PtLhcx1 | Lhcx1 PHATRDRAFT_27278 | XP_002179760.1; | Protein fucoxanthin chlorophyll a/c protein |
| PtLhcr9 | Lhcr9 PHATR_43860 | XP_002186024.1; | Fucoxanthin chlorophyll a/c protein, lhcr type |
| PtLhcr8 | Lhcr8 PHATRDRAFT_32294 | XP_002176917.1; | Protein fucoxanthin chlorophyll a/c protein |
| PtLhcr7 | Lhcr7 PHATRDRAFT_43522 | XP_002177668.1; | Protein fucoxanthin chlorophyll a/c protein |
| PtLhcr6 | Lhcr6 PHATRDRAFT_56319 | XP_002181976.1; | Protein fucoxanthin chlorophyll a/c protein |
| PtLhcr5 | Lhcr5 PHATRDRAFT_29472 | XP_002182761.1; | Protein fucoxanthin chlorophyll a/c protein (Fragment) |
| PtLhcr4 | Lhcr4 PHATRDRAFT_17766 | XP_002177385.1; | Protein fucoxanthin chlorophyll a/c protein |
| PtLhcr3 | Lhcr3 PHATRDRAFT_50725 | XP_002178019.1; | Protein fucoxanthin chlorophyll a/c protein |
| PtLhcr2 | Lhcr2 PHATRDRAFT_22956 | XP_002183608.1; | Protein fucoxanthin chlorophyll a/c protein |
| PtLhcr14 | Lhcr14 PHATRDRAFT_47813 | XP_002182162.1; | Protein fucoxanthin chlorophyll a/c protein |
| PtLhcr13 | Lhcr13 PHATRDRAFT_38121 | XP_002182329.1; | Protein fucoxanthin chlorophyll a/c protein |
| PtLhcr12 | Lhcr12 PHATRDRAFT_54027 | XP_002176857.1; | Protein fucoxanthin chlorophyll a/c protein |
| PtLhcr11 | Lhcr11 PHATRDRAFT_23257 | XP_002184127.1; | Protein fucoxanthin chl a/c protein |
| PtLhcr10 | Lhcr10 PHATRDRAFT_50086 | XP_002184869.1; | Protein fucoxanthin chlorophyll a/c protein |
| PtLhcr1 | Lhcr1 PHATRDRAFT_44601 | XP_002178624.1; | Protein fucoxanthin chlorophyll a/c protein |
| PtLhcf9 | Lhcf9 PHATRDRAFT_30031 | XP_002183709.1; | Protein fucoxanthin chlorophyll a/c protein |
| PtLhcf8 | Lhcf8 PHATRDRAFT_22395 | XP_002182937.1; | Protein fucoxanthin chlorophyll a/c protein |
| PtLhcf6 | Lhcf6 Lhcf7 PHATRDRAFT_29266 PHATRDRAFT_30643 | XP_002182305.1; | Protein fucoxanthin chlorophyll a/c protein |
| PtLhcf7 | | XP_002184540.1; | |
| PtLhcf5 | Lhcf5 PHATRDRAFT_30648 | XP_002184620.1; | Protein fucoxanthin chlorophyll a/c protein |
| PtLhcf4 | Lhcf4 Lhcf3 PHATRDRAFT_25168 PHATRDRAFT_50705 | XP_002177868.1; | Protein fucoxanthin chlorophyl a/c protein (Protein fucoxanthin chlorophyll a/c protein) |
| PtLhcf3 | | XP_002177869.1; | |
| PtLhcf2 | Lhcf2 PHATRDRAFT_25172 | XP_002177870.1; | Protein fucoxanthin chlorophyll a/c protein |
| PtLhcf17 | Lhcf17 PHATRDRAFT_56310 | XP_002184763.1; | Protein fucoxanthin chlorophyll a/c protein |
| **PtLhcq4** | Lhcf16 PHATRDRAFT_34536 | XP_002178860.1; | Protein fucoxanthin chlorophyll a/c protein |
| PtLhcf15 | Lhcf15 PHATRDRAFT_48882 | XP_002183381.1; | Protein fucoxanthin chlorophyll a/c protein |
| PtLhcf14 | Lhcf14 PHATR_25893 | XP_002186206.1; | Fucoxanthin chlorophyll a/c protein, lhcf type |
| PtLhcf13 | Lhcf13 PHATRDRAFT_22680 | XP_002183291.1; | Protein fucoxanthin chlorophyll a/c protein |
| PtLhcf12 | Lhcf12 PHATRDRAFT_16302 | XP_002184765.1; | Protein fucoxanthin chlorophyll protein |
| PtLhcf11 | Lhcf11 PHATRDRAFT_51230 | XP_002184619.1; | Protein fucoxanthin chlorophyll a/c protein |
| PtLhcf10 | Lhcf10 PHATRDRAFT_22006 | XP_002182219.1; | Protein fucoxanthin chlorophyll a/c protein |
| PtLhcf1 | Lhcf1 PHATRDRAFT_18049 | XP_002177871.1; | Protein fucoxanthin chlorophyll a/c protein |

### Supplemental Table S5. List of RefSeq or GenBank accession IDs or other references used to obtain the FCP/LHC sequences.

| Lineage | Symbiosis | Order | RefSeq/GenBank or Other Assembly Accession | Species |
|---|---|---|---|---|
| Red | 1st | Rhodophyta; Bangiophyceae; Cyanidiales; Cyanidiaceae | GCF_000091205.1 | *Cyanidioschyzon merolae* |
| Red | 1st | Rhodophyta; Bangiophyceae; Porphyridiales; Porphyridiaceae | GCA_008690995.1 | *Porphyridium purpureum* |
| Red | 2nd | Stramenopiles; Ochrophyta; PX clade; Phaeophyceae; Ectocarpales; Ectocarpaceae | GCA_000310025.1 | *Ectocarpus siliculosus* |
| Red | 2nd | Stramenopiles; Ochrophyta; Raphidophyceae; Chattonellales; Chattonellaceae | Harmful Algal Blooms (DB-HABs, http://hab.nibb.ac.jp/) | *Chattonella antiqua* |
| Red | 3rd | Alveolata; Dinophyceae; Peridiniales; Heterocapsaceae | Harmful Algal Blooms (DB-HABs, http://hab.nibb.ac.jp/) | *Heterocapsa circularisquama* |
| Red | 2nd | Stramenopiles; Ochrophyta; Bacillariophyta; Bacillariophyceae; Bacillariophycidae; Bacillariales; Bacillariaceae | GCA_900660405.1 | *Pseudo-nitzschia multistriata* |
| Red | 2nd | Stramenopiles; Ochrophyta; Bacillariophyta; Bacillariophyceae; Bacillariophycidae; Bacillariales; Bacillariaceae | GCA_001750085.1 | *Fragilariopsis cylindrus* |
| Red | 2nd | Stramenopiles; Ochrophyta; Bacillariophyta; Bacillariophyceae; Bacillariophycidae; Naviculales; Naviculaceae | GCA_002217885.1 | *Fistulifera solaris* |
| Red | 2nd | Stramenopiles; Ochrophyta; Bacillariophyta; Bacillariophyceae; Bacillariophycidae; Naviculales; Phaeodactylaceae | GCF_000150955.2 | *Phaeodactylum tricornutum* |
| Red | 2nd | Stramenopiles; Ochrophyta; Bacillariophyta; Coscinodiscophyceae; Chaetocerotophycidae; Chaetocerotales; Chaetocerotaceae | | *Chaetoceros gracilis* |
| Red | 2nd | Stramenopiles; Ochrophyta; Bacillariophyta; Coscinodiscophyceae; Thalassiosirophycidae; Thalassiosirales; Thalassiosiraceae | GCF_000149405.2 | *Thalassiosira pseudonana* |
| Red | 2nd | Stramenopiles; Ochrophyta; Bacillariophyta; Coscinodiscophyceae; Thalassiosirophycidae; Thalassiosirales; Thalassiosiraceae | GCA_000296195.2 | *Thalassiosira oceanica* |
| Red | 2nd | Haptista; Haptophyta; Prymnesiophyceae; Isochrysidales; Noelaerhabdaceae | GCF_000372725.1 | *Emiliania huxleyi* |
| Red | 2nd | Haptista; Haptophyta; Prymnesiophyceae; Prymnesiales; Chrysochromulinaceae | GCA_001275005.1 | *Chrysochromulina tobinii* |
| Red | 2nd | Haptista; Haptophyta; Prymnesiophyceae; Phaeocystales; Phaeocystaceae | Phaeocystis antarctica CCMP1374 v2.2, Proposal ID: 869 / 99187, https://genome.jgi.doe.gov/portal/Phaant1/Phaant1.download.html | *Phaeocystis antarctica* |
| Green | 1st | Viridiplantae; Chlorophyta; core chlorophytes; Chlorophyceae; Chlamydomonadales; Chlamydomonadaceae | GCF_000002595.1 | *Chlamydomonas reinhardtii* |
| Green | 1st | Viridiplantae; Streptophyta; Streptophytina; Embryophyta; Bryophyta; Bryophytina; Bryopsida; Funariidae; Funariales; Funariaceae | GCF_000002425.4 | *Physcomitrella patens* |
| Red | NA | Cryptophyceae; Pyrenomonadales; Geminigeraceae | GCF_000315625.1 | *Guillardia theta* |

**Supplemental Table S6. The conserved FCP set of diatoms, including the FCPs assigned to *Chaetoceros gracilis* photosystems.** FCPs of other than *Chaetoceros gracilis* and two model diatoms were noted as GenBank accession ID. FCPs of *Chaetoceros gracilis* and two model diatoms were noted as gene names listed in Supplemental table S3 and S4. Gene names with bold letters are renamed from former annotations.

| *C. gracilis* | *T. pseudonana* | *T. oceanica* | *F. cylindrus* | *F. solaris* | *P. tricornutum* |
|---|---|---|---|---|---|
| CgLhcr1 | TpLhcr3 | EJK62427.1 | OEU06127.1, OEU13720.1 | GAX12308.1 | PtLhcr3 |
| CgLhcr2, CgLhcr6 | TpLhcr4, (TpLhcr14) | EJK72728.1, EJK72727.1 | OEU16552.1, OEU16551.1 | GAX21829.1, GAX13743.1, (GAX21830.1, GAX13742.1) | PtLhcr4, (PtLhcr12) |
| CgLhcr3 | **TpLhcr18** | EJK71517.1 | OEU13749.1 | GAX13128.1 | PtLhcr14 |
| CgLhcr4 | **TpLhcq7** | EJK45846.1 | OEU21089.1 | GAX28645.1, GAX19964.1 | **PtLhcq2** |
| CgLhcr5 | TpLhcr1 | EJK51776.1, EJK46172.1 | OEU18936.1 | GAX16008.1, GAX11451.1 | PtLhcr1 |
| CgLhcr7 | TpLhcr7, **TpLhcr19** | EJK49415.1 | OEU14130.1 | GAX24797.1 | PtLhcr2 |
| CgLhcr8 | TpLhcr11, TpLhcr12 | EJK77608.1 | OEU12359.1 | GAX09492.1 | PtLhcr11 |
| CgLhcr9 | **TpLhcq10** | EJK50038.1 | OEU11937.1 | GAX14579.1 | **PtLhcq5** |
| CgLhcr10 | **TpLhcr20** | EJK71515.1 | OEU13748.1 | GAX13127.1 | PtLhcr13 |
| CgLhcr17 | **TpLhcr17** | EJK46744.1 | OEU09133.1 | GAX18319.1, GAX16656.1 | **PtLhcr16** |

### Supplemental Table S7. List of RefSeq or GenBank accession IDs used to infer phylogenetic tree of chloroplast genes.

| Taxon | Species | Accession Refseq/GenBank | Note |
|---|---|---|---|
| Alveolata; Dinophyceae; Gymnodiniales; Gymnodiniaceae | *Lepidodinium chlorophorum* | NC_027093.1 | |
| Alveolata; Dinophyceae; Peridiniales; Kryptoperidiniaceae | *Durinskia baltica* | NC_014287.1 | |
| Alveolata; Dinophyceae; Peridiniales; Kryptoperidiniaceae | *Kryptoperidinium foliaceum* | NC_014267.1 | |
| Cryptophyceae; Cryptomonadales; Cryptomonadaceae | *Cryptomonas curvata* | NC_035720.1 | |
| Cryptophyceae; Pyrenomonadales; Pyrenomonadaceae | *Rhodomonas salina* | NC_009573.1 | |
| Glaucocystophyceae; Cyanophoraceae | *Cyanophora paradoxa* | NC_001675.1 | *cyanelle |
| Haptista; Haptophyta; Prymnesiophyceae; Isochrysidales; Isochrysidaceae | *Isochrysis galbana* | NC_049168.1 | |
| Haptista; Haptophyta; Prymnesiophyceae; Isochrysidales; Noelaerhabdaceae | *Emiliania huxleyi* | NC_007288.1 | |
| Haptista; Haptophyta; Prymnesiophyceae; Prymnesiales; Chrysochromulinaceae | *Chrysochromulina parva* | NC_036937.1 | |
| Rhodophyta; Bangiophyceae; Bangiales; Bangiaceae | *Porphyra umbilicalis* | NC_035573.1 | |
| Rhodophyta; Bangiophyceae; Cyanidiales; Cyanidiaceae | *Cyanidioschyzon merolae* | NC_004799.1 | |
| Rhodophyta; Bangiophyceae; Cyanidiales; Cyanidiaceae | *Galdieria sulphuraria* | NC_024665.1 | |
| Rhodophyta; Bangiophyceae; Porphyridiales; Porphyridiaceae | *Porphyridium purpureum* | NC_023133.1 | |
| Stramenopiles; Ochrophyta; Bacillariophyta; Bacillariophyceae; Bacillariophycidae; Naviculales; Phaeodactylaceae | *Phaeodactylum tricornutum* | NC_008588.1 | |
| Stramenopiles; Ochrophyta; Bacillariophyta; Coscinodiscophyceae; Chaetocerotophycidae; Chaetocerotales; Chaetocerotaceae | *Chaetoceros simplex* | NC_025310.1 | |
| Stramenopiles; Ochrophyta; Bacillariophyta; Coscinodiscophyceae; Thalassiosirophycidae; Thalassiosirales; Thalassiosiraceae | *Thalassiosira pseudonana* | NC_008589.1 | |
| Stramenopiles; Ochrophyta; Bolidophyceae; Parmales; Triparmaceae | *Triparma laevis* | NC_027746.1 | |
| Stramenopiles; Ochrophyta; Dictyochophyceae; Dictyochales | *Dictyocha speculum* | NC_043929.1 | |
| Stramenopiles; Ochrophyta; Dictyochophyceae; Rhizochromulinales | *Rhizochromulina marina* | NC_043890.1 | |
| Stramenopiles; Ochrophyta; Eustigmatophyceae | *Nannochloropsis oculata* | NC_022260.1 | |
| Stramenopiles; Ochrophyta; PX clade; Phaeophyceae; Dictyotales; Dictyotaceae | *Dictyopteris divaricata* | KY433579.1 | |
| Stramenopiles; Ochrophyta; PX clade; Phaeophyceae; Ectocarpales; Ectocarpaceae | *Ectocarpus siliculosus* | FP102296.1 | |
| Stramenopiles; Ochrophyta; PX clade; Xanthophyceae | *Vaucheria litorea* | NC_011600.1 | |
| Stramenopiles; Ochrophyta; Raphidophyceae; Chattonellales; Chattonellaceae | *Heterosigma akashiwo* | NC_010772.1 | |
| Stramenopiles; Pelagophyceae; Pelagomonadales | *Aureoumbra lagunensis* | NC_012903.1 | |
| Stramenopiles; Pelagophyceae; Pelagomonadales | *Aureococcus anophagefferens* | NC_012898.1 | |
| Viridiplantae; Chlorophyta; core chlorophytes; Chlorophyceae; Chlamydomonadales; Chlamydomonadaceae | *Chlamydomonas reinhardtii* | NC_005353.1 | |
| Viridiplantae; Chlorophyta; core chlorophytes; Chlorophyceae; Chlamydomonadales; Volvocaceae | *Volvox africanus* | NC_039755.1 | |
